# Expression quantitative trait loci-derived scores and white matter microstructure in UK Biobank: a novel approach to integrating genetics and neuroimaging

**DOI:** 10.1101/646646

**Authors:** Miruna C. Barbu, Athina Spiliopoulou, Marco Colombo, Paul McKeigue, Toni-Kim Clarke, David M. Howard, Mark J. Adams, Xueyi Shen, Stephen M. Lawrie, Andrew M. McIntosh, Heather C. Whalley

## Abstract

**Background:** Expression quantitative trait loci (eQTL) are genetic variants associated with gene expression. Using genome-wide genotype data, it is now possible to impute gene expression using eQTL mapping efforts. This approach can be used to analyse previously unexplored relationships between gene expression and heritable *in vivo* measures of human brain structural connectivity.

**Methods:** Using large-scale eQTL mapping studies, we computed 6,457 gene expression scores (eQTL scores) using genome-wide genotype data in UK Biobank, where each score represents a genetic proxy measure of gene expression. These scores were then tested for associations with two diffusion tensor imaging measures, fractional anisotropy (N_FA_=14,518) and mean diffusivity (N_MD_=14,485), representing white matter structural integrity.

**Results:** We found FDR-corrected significant associations between 8 eQTL scores and structural connectivity phenotypes, including global and regional measures (β_absolute_ FA=0.0339-0.0453; MD=0.0308-0.0381) and individual tracts (β_absolute_ FA=0.0320-0.0561; MD=0.0295-0.0480). The loci within these eQTL scores have been reported to regulate expression of genes involved in various brain-related processes and disorders, such as neurite outgrowth and Parkinson’s disease (*DCAKD*, *SLC35A4*, *SEC14L4*, *SRA1*, *NMT1*, *CPNE1*, *PLEKHM1*, *UBE3C*).

**Discussion:** Our findings indicate that eQTL scores are associated with measures of *in vivo* brain connectivity and provide novel information not previously found by conventional genome-wide association studies. Although the role of expression of these genes regarding white matter microstructural integrity is not yet clear, these findings suggest it may be possible, in future, to map potential trait- and disease-associated eQTL to *in vivo* brain connectivity and better understand the mechanisms of psychiatric disorders and brain traits, and their associated imaging findings.

## Introduction

Expression quantitative trait loci (eQTL) are genetic variants which are proximally (cis) or distally (trans) associated with variation in the expression of genes (1). Previous animal and human studies have found that changes in gene expression lead to phenotypic variation, including adaptive phenotypic changes and evolutionary developments. In humans, for instance, cis-regulatory mutations lead to differences in lactase (*LCT*) gene expression, resulting in lactase persistence in adulthood (2). With respect to psychiatric disorders, major depressive disorder (MDD) and bipolar disorder have been associated with decreased expression of prodynorphin messenger RNA (mRNA), which is involved in regulation of mood and expressed in limbic-related areas within the brain (e.g. amygdala, hippocampus) (3; 4; 5). These findings indicate the importance of cis-regulatory mutations and variations in trait evolution.

Variation in gene regulation leads to differences in individual phenotypes, indicating that eQTL may play a role in susceptibility to disease (6; 7). To test this hypothesis, methods which combine gene expression data with genome-wide association studies (GWAS) summary statistics have been developed. These approaches may provide further insight into the potential causal pathways and genes involved in specific disorders, or predict the regulatory roles of single nucleotide polymorphisms (SNPs) in linkage disequilibrium (LD) with previously associated variants (8). Previous studies have found that genetic variation may explain some of the variance in levels of gene expression in human tissues, including post-mortem brain tissue (9; 10; 11; 12). In one such study, Zou et al. (2012) (13) conducted an expression genome-wide association study (eGWAS) on post-mortem brains of individuals with Alzheimer’s disease (AD) and other brain pathologies (non-AD; including progressive supranuclear palsy). They found 2,980 *cis*SNPs associated with both AD and non-AD conditions. By investigating brain eQTL in post-mortem tissue therefore, researchers have been able to discover associations between gene expression and disease states in the brain.

Using brain tissue in order to investigate gene expression levels is however problematic, due to limitations such as small sample sizes and possible expression level differences in post-mortem versus ante-mortem brains (14). As such, alternative approaches have therefore been investigated. One such approach is using eQTL measured from whole blood gene expression as a proxy for brain gene expression; an approach supported by important benefits such as greater sample size and easier accessibility (15). Although it is recommended that wherever possible gene expression levels should be measured in a tissue-specific manner, considerable overlap has been demonstrated between blood and brain eQTL, indicating the validity of the approach (14).

Neuroimaging measures provide a novel opportunity to investigate whether eQTL are significantly associated with *in vivo* brain phenotypes, and thereby increasing our knowledge of the role of eQTL in the wider context of psychiatric disorders. White matter microstructure, as measured by diffusion tensor imaging (DTI), is consistently heritable across tracts (16; 17; 18) and is compromised in several psychiatric disorders. Generally, decreased microstructural integrity of white matter is characterised by lower directionality of water molecule diffusion (reduced fractional anisotropy, FA) and less constrained water molecule diffusion (increased mean diffusivity, MD). Consistent findings across studies have indicated higher MD and lower FA in individuals suffering from MDD, for example (19; 20). Investigating the regulatory loci associated with white matter microstructure in health and disease may aid in the detection of molecular mechanisms influencing disease through aberrant structural brain connectivity.

Within the current study, we derived eQTL scores based on two well-powered whole-blood eQTL studies (21; 22). We then used GENOSCORES, a database of filtered summary statistics of publicly-available GWAS covering multiple phenotypes, including gene expression, to calculate eQTL scores (https://pm2.phs.ed.ac.uk/genoscores/).

The resultant eQTL-based genetic scores can be considered proxies for the expression of particular genes, which can then be tested for association with traits of interest. Here, we analysed their association with white matter microstructure as measured by FA and MD in UK Biobank. We used participants from the October 2018 UK Biobank neuroimaging release (N_FA_ = 14,518; N_MD_ = 14,485). The purpose of the study was to utilise a novel approach to investigate associations between regulatory SNPs and white matter microstructure. This approach could lead to further specialised investigation into psychiatric and neurological disorders, as well as other brain-related traits, such as cognition and behaviour.

## Methods and materials

### UK Biobank (UKB)

UK Biobank is a health resource aiming to prevent, diagnose and treat numerous disorders. It is comprised of 502,617 individuals whose genetic and environmental data (e.g. lifestyle, medications) were collected between 2006 and 2010 in the United Kingdom (http://www.ukbiobank.ac.uk/). UKB received ethical approval from the Research Ethics Committee (reference: 11/NW/0382). This study has been approved by the UKB Access Committee (Project #4844). Written informed consent was obtained from all participants.

### Study population

In the current study, individuals were excluded if they participated in studies such as the Psychiatric Genomics Consortium (PGC) MDD GWAS or Generation Scotland (Scottish Family Health Study) to remove overlap of genetic samples. For the brain imaging sample, a quality check performed by UK Biobank ensured that no abnormal scans were included in subsequent analyses (23). We additionally excluded individuals whose global measures for FA and MD lay more than three standard deviations from the sample mean (19; 24). This resulted in 14,518 individuals with FA values (N_female_ = 7,561 (52%); N_male_ = 6,957 (48%); mean age: 63.14 +/− 7.4; age range: 45.92 – 80.67) and 14,485 individuals with MD values (N_female_ = 7,552 (52%); N_male_ = 6,933 (48%); mean age: 63.12 +/− 7.39; age range: 45.92 – 80.67).

### Genotyping and eQTL score calculation

A total of 488,363 UKB blood samples (N female = 264,857; N male = 223,506; http://biobank.ctsu.ox.ac.uk/crystal/field.cgi?id=22001) were genotyped using the UK BiLEVE array (N = 49,949; http://biobank.ctsu.ox.ac.uk/crystal/refer.cgi?id=149600) and the UK Biobank Axiom array (N = 438,417; http://biobank.ctsu.ox.ac.uk/crystal/refer.cgi?id=149601). Details of genotyping and quality control are described in more detail by Hagenaars et al. (2016) (25) and Bycroft et al. (2017) (26).

From GENOSCORES, we used eQTL analysis summary statistics from two studies of whole-blood eQTL (21; 22). Briefly, Gusev et al. (2016) (21) developed a novel approach aimed at identifying associations between gene expression and complex traits in cases where gene expression level is not directly measured. These authors reported eQTL based on a sample of 1,414 individuals with whole-blood expression measured using the Illumina HumanHT-12 version 4 Expression BeadChip. Westra et al. (2013) (22) performed a large eQTL meta-analysis in 5,311 samples across 7 studies from peripheral blood, with gene expression measured using Illumina whole-genome Expression BeadChips (HT12v3, HT12v4 or H8v2 arrays). Their aim was to investigate the magnitude of the effect of cis and trans SNPs on gene expression, as well as to observe whether mapping eQTL in peripheral blood could uncover biological pathways associated with complex traits and disease. Further details of data acquisition and protocols are described in more detail in the two studies (21; 22).

Before being imported into the GENOSCORES database, summary statistics were filtered at a liberal p-value < 1E-4 (0.0001). We computed a total of 10,884 eQTL scores (N Gusev study = 3,801; N Westra study = 7,083) for individuals included in the imaging sample (N_FA_: 14,518; N_MD_: 14,485) from the SNPs found in GENOSCORES, using a p-value threshold of 1E-5 (0.00001). We then excluded overlapping eQTL scores between the two studies (i.e. scores for which SNPs affect expression of the same gene in both studies) by only including the score where a SNP had the lowest p-value, i.e. most significant association. The final eQTL score list was 6,457 (N Gusev study = 3,286; N Westra study = 3,171). These scores were used as input variables in subsequent statistical analyses. Figure 1 below provides a summary of the score derivation process.

**Figure 1.**
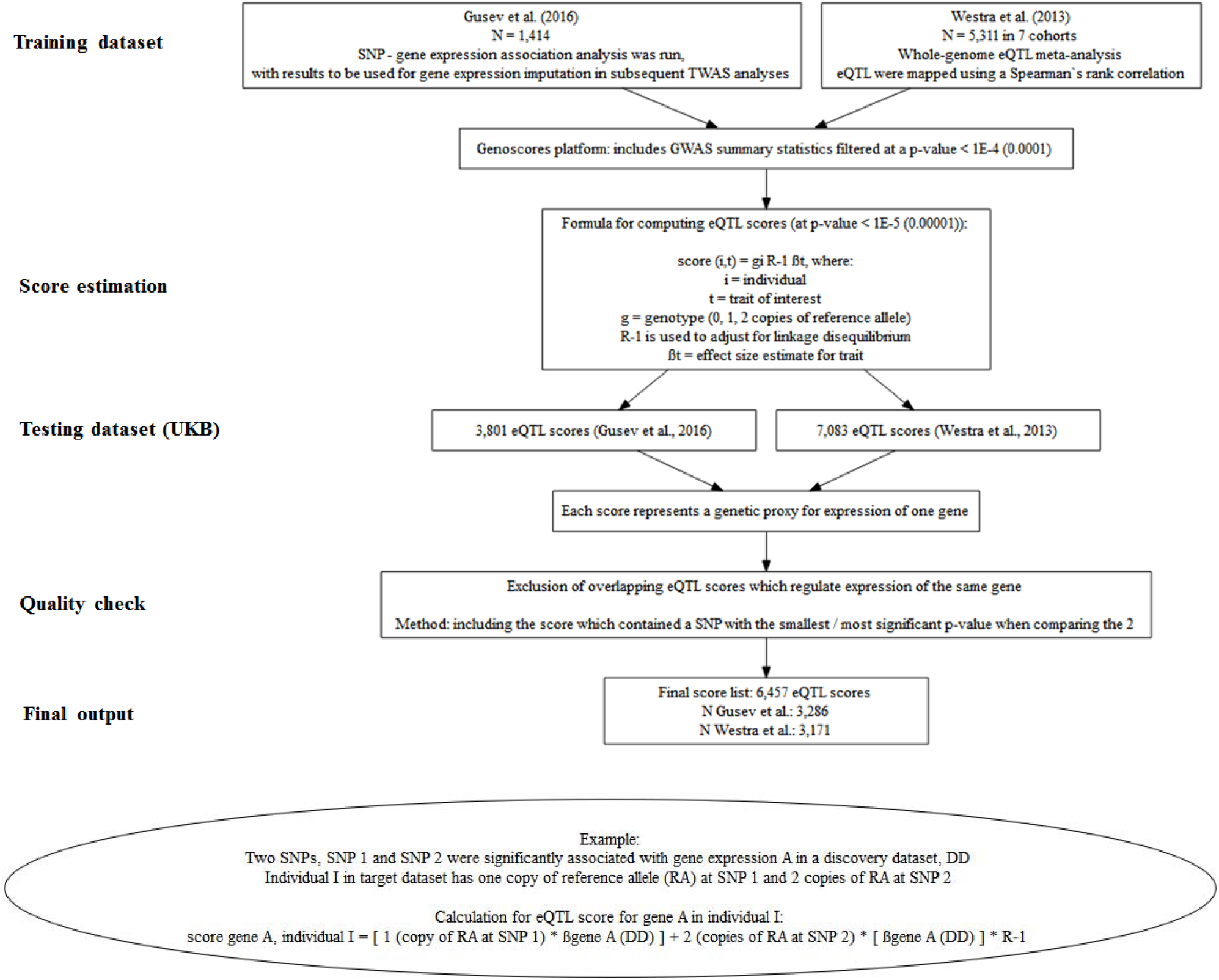
eQTL score computation process.

Briefly, eQTL scores were computed as a sum of the genotypes for an individual (g, scored as 0, 1, 2 copies of the reference allele) weighted by the effect size estimate (*βt*) for the trait of interest *t.* In order to adjust for LD, vector *βt* was pre-multiplied by the generalized inverse of the SNP-SNP correlation matrix R estimated from the 1000 Genomes reference panel, limited to the individuals with European ancestry.

The formula to compute the eQTL score for trait *t* for an individual (*i*) is therefore:

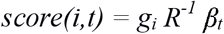

### Magnetic resonance imaging (MRI) acquisition

In the current study, imaging-derived phenotypes (IDPs) produced by UKB were used. MRI acquisition and pre-processing procedures for white matter tracts were performed by UKB using standardised protocols (https://biobank.ctsu.ox.ac.uk/crystal/docs/brain_mri.pdf). Briefly, images were acquired in Manchester (N_FA_ = 12,248; N_MD_ = 12,221) and Newcastle (N_FA_ = 2,270; N_MD_ = 2,264) on a standard Siemens Skyra 3T scanner with a 32-channel radio-frequency (RF) receive head coil and later pre-processed using the FMRIB Software Library (FSL), and parcellation of white matter tracts was conducted using AutoPtx (23). Individual white matter tracts belonging to each tract category can be observed in supplementary materials section 5.

Owing to the fact that head position and RF coil in the scanner may affect data quality and subsequent pre-processing, three scanner brain position variables were also generated by UKB, with the aim of being used as confounding variables in subsequent analyses. These are lateral brain position – X (http://biobank.ctsu.ox.ac.uk/crystal/field.cgi?id=25756), transverse brain position – Y (http://biobank.ctsu.ox.ac.uk/crystal/field.cgi?id=25757) and longitudinal brain position – Z (http://biobank.ctsu.ox.ac.uk/crystal/field.cgi?id=25758). The three variables were included as covariates in the statistical analysis described below.

### Statistical methods

All analyses were conducted using R (version 3.2.3) in a Linux environment. For each white matter tract, we used generalized linear mixed models (function “lme” in package “nlme”) for bilateral brain regions, which were included as dependent variables. The eQTL scores were included as independent variables separately in each model, with additional covariates: age, age^2^, sex, fifteen genetic principal components to control for population stratification, three MRI head position coordinates, MRI site and genotype array, while hemisphere was included as a within-subject variable. For unilateral tracts, as well as global measures and white matter tract categories of FA and MD, also included in the models as dependent variables, we used a general linear model (function “lm”), using the same covariates as above, without hemisphere included as a separate term, and again including the eQTL scores as independent variables separately in each model.

For global measures and white matter tract categories of FA and MD, we applied principal component analysis (PCA) on the white matter tracts of interest (all 27 for global measures; 12 for association fibres; 6 for thalamic radiations; 9 for projection fibres) in order to extract a latent measure. Scores of the first unrotated component were extracted and set as dependent variables in general linear models. False discovery rate (FDR) correction using the “p.adjust” function in R (q < 0.05) was applied across the eQTL scores and the individual white matter tracts (N_tests_ = 98,855), and separately across eQTL scores and global and tract categories (N_tests_ = 25,828).

## Results

There were several eQTL scores that showed significant associations with a number of global measures, tract categories, and white matter tracts post FDR correction (Table 1; Figures 2a & 3b and 3a & 3b; supplementary section 3). In total, 25 scores were significantly associated with FA values (β_absolute_ = 0.0320-0.0561) and 24 scores with MD values (β_absolute_ = 0.0295-0.0480) in several tracts (see supplementary material section 1). Among these scores, 8 were associated with white matter tracts measured by both FA and MD. The primary findings reported in this manuscript focus on these 8 overlapping scores, as these were considered to provide the most consistent information with regards to gene expression within white matter tracts as measured by two different DTI scalars (see tables 2–7 below), further findings are presented in supplementary materials.

**Table 1.** Information regarding eQTL scores with significant associations for both FA and MD-measured tracts.

| Score name & eQTL type | N SNPs in score | Regulated gene | Study from which score is calculated | Gene function |
| --- | --- | --- | --- | --- |
| DCAKD_eQTL_cis | 8 | DCAKD | Gusev et al. | Expressed in glioma; ubiquitous expression in brain; implicated in a number of psychiatric and neurological disorders (31; 32; 33; 34) |
| SLC35A4_eQTL_cis | 12 | SLC35A4 | Gusev et al. | Expressed in brain (36) |
| SEC14L4_eQTL_cis | 1 | SEC14L4 | Westra et al. | Specific function not yet determined; may be implicated in neurodegeneration (41) |
| SRA1_eQTL_cis | 15 | SRA1 | Westra et al. | Involved in regulation of many NR (nuclear receptor) and non-NR activities (e.g. chromatin organisation); may be associated with idiopathic hypogonadotropic hypogonadism (37; 38) |
| NMT1_eQTL_cis | 7 | NMT1 | Westra et al. | Ubiquitous expression in brain; may be implicated in brain tumours (47; 48; 49) |
| CPNE1_eQTL_cis | 1 | CPNE1 | Westra et al. | May regulate molecular events at the interface of the cell membrane and cytoplasm; expressed during brain development and implicated in neurite outgrowth in rats (44; 45; 46) |
| PLEKHM1_eQTL_cis | 5 | PLEKHM1 | Gusev et al. | Protein encoded by this gene is important for bone resorption; may play critical role in vesicular transport in the osteoclast (42; 43) |
| UBE3C_eQTL_cis | 4 | UBE3C | Westra et al. | Expressed in brain; may be implicated in Parkinson's disease (39; 40) |

### The effect of the 8 scores on FA measures of white matter microstructure

**Table 2.** Significant associations between eQTL scores and global FA; FDR = false discovery rate.

| Score, White Matter Tracts | Effect size | SD | t value | p value | p value, FDR corrected |
| --- | --- | --- | --- | --- | --- |
| **DCAKD eQTL score** |  |  |  |  |  |
| Global FA | -0.0367 | 0.0079 | -4.6474 | 3.39161E-06 | 0.0088 |
| **SLC35A4 eQTL score** |  |  |  |  |  |
| Global FA | -0.0403 | 0.0079 | -5.0996 | 3.44595E-07 | 0.0022 |
| **SEC14L4 eQTL score** |  |  |  |  |  |
| Global FA | -0.0420 | 0.0079 | -5.3199 | 1.0538E-07 | 0.0011 |

**Table 3.** Significant associations between eQTL scores and tract categories (FA); FDR = false discovery rate; Effect size in table is ordered from smallest to largest for each score.

| Score, White Matter Tracts | Effect size | SD | t value | p value | p value, FDR corrected |
| --- | --- | --- | --- | --- | --- |
| **DCAKD eQTL score** |  |  |  |  |  |
| Thalamic radiations | -0.0403 | 0.0080 | -5.0378 | 4.76577E-07 | 0.0025 |
| **SLC35A4 eQTL score** |  |  |  |  |  |
| Association fibres | -0.0347 | 0.0079 | -4.4036 | 1.07241E-05 | 0.0198 |
| Projection fibres | -0.0453 | 0.0079 | 5.7612 | 8.51978E-09 | 0.0002 |
| **SEC14L4 eQTL score** |  |  |  |  |  |
| Association fibres | -0.0358 | 0.0079 | -4.5425 | 5.6047E-06 | 0.0121 |
| Thalamic radiations | -0.0388 | 0.0080 | -4.8429 | 1.2928E-06 | 0.0048 |
| Projection fibres | -0.0416 | 0.0079 | 5.2850 | 1.275E-07 | 0.0011 |
| **SRA1 eQTL score** |  |  |  |  |  |
| Projection fibres | -0.0339 | 0.0079 | 4.3032 | 1.6943E-05 | 0.0273 |

**Table 4.**
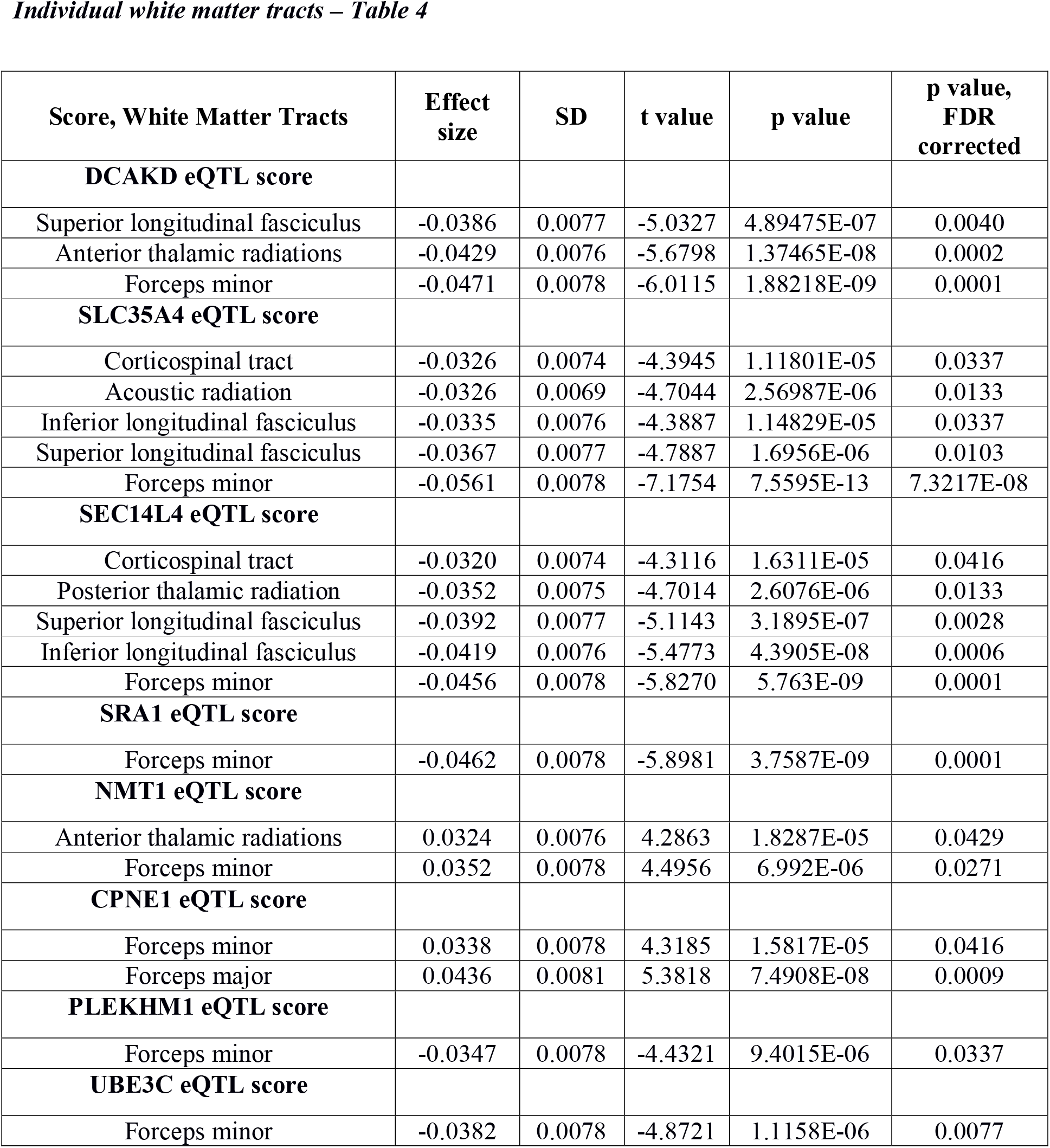
Significant associations between eQTL scores and individual white matter tracts (FA); FDR = false discovery rate; Effect size in table is ordered from smallest to largest for each score.

| Score, White Matter Tracts | Effect size | SD | t value | p value | p value, FDR corrected |
| --- | --- | --- | --- | --- | --- |
| **DCAKD eQTL score** |  |  |  |  |  |
| Superior longitudinal fasciculus | -0.0386 | 0.0077 | -5.0327 | 4.89475E-07 | 0.0040 |
| Anterior thalamic radiations | -0.0429 | 0.0076 | -5.6798 | 1.37465E-08 | 0.0002 |
| Forceps minor | -0.0471 | 0.0078 | -6.0115 | 1.88218E-09 | 0.0001 |
| **SLC35A4 eQTL score** |  |  |  |  |  |
| Corticospinal tract | -0.0326 | 0.0074 | -4.3945 | 1.11801E-05 | 0.0337 |
| Acoustic radiation | -0.0326 | 0.0069 | -4.7044 | 2.56987E-06 | 0.0133 |
| Inferior longitudinal fasciculus | -0.0335 | 0.0076 | -4.3887 | 1.14829E-05 | 0.0337 |
| Superior longitudinal fasciculus | -0.0367 | 0.0077 | -4.7887 | 1.6956E-06 | 0.0103 |
| Forceps minor | -0.0561 | 0.0078 | -7.1754 | 7.5595E-13 | 7.3217E-08 |
| **SEC14L4 eQTL score** |  |  |  |  |  |
| Corticospinal tract | -0.0320 | 0.0074 | -4.3116 | 1.6311E-05 | 0.0416 |
| Posterior thalamic radiation | -0.0352 | 0.0075 | -4.7014 | 2.6076E-06 | 0.0133 |
| Superior longitudinal fasciculus | -0.0392 | 0.0077 | -5.1143 | 3.1895E-07 | 0.0028 |
| Inferior longitudinal fasciculus | -0.0419 | 0.0076 | -5.4773 | 4.3905E-08 | 0.0006 |
| Forceps minor | -0.0456 | 0.0078 | -5.8270 | 5.763E-09 | 0.0001 |
| **SRA1 eQTL score** |  |  |  |  |  |
| Forceps minor | -0.0462 | 0.0078 | -5.8981 | 3.7587E-09 | 0.0001 |
| **NMT1 eQTL score** |  |  |  |  |  |
| Anterior thalamic radiations | 0.0324 | 0.0076 | 4.2863 | 1.8287E-05 | 0.0429 |
| Forceps minor | 0.0352 | 0.0078 | 4.4956 | 6.992E-06 | 0.0271 |
| **CPNE1 eQTL score** |  |  |  |  |  |
| Forceps minor | 0.0338 | 0.0078 | 4.3185 | 1.5817E-05 | 0.0416 |
| Forceps major | 0.0436 | 0.0081 | 5.3818 | 7.4908E-08 | 0.0009 |
| **PLEKHM1 eQTL score** |  |  |  |  |  |
| Forceps minor | -0.0347 | 0.0078 | -4.4321 | 9.4015E-06 | 0.0337 |
| **UBE3C eQTL score** |  |  |  |  |  |
| Forceps minor | -0.0382 | 0.0078 | -4.8721 | 1.1158E-06 | 0.0077 |

### The effect of the 8 scores on MD measures of white matter microstructure

**Table 5.** Significant associations between eQTL scores and global MD; FDR = false discovery rate.

| Score, White Matter Tracts | Effect size | SD | t value | p value | p value, FDR corrected |
| --- | --- | --- | --- | --- | --- |
| **DCAKD eQTL score** |  |  |  |  |  |
| Global MD | 0.0404 | 0.0075 | 5.3762 | 7.72382E-08 | 0.0015 |
| **SLC35A4 eQTL score** |  |  |  |  |  |
| Global MD | 0.0308 | 0.0075 | 4.0893 | 4.3502E-05 | 0.0423 |
| **SEC14L4 eQTL score** |  |  |  |  |  |
| Global MD | 0.0326 | 0.0075 | 4.3299 | 1.5015E-05 | 0.0277 |
| **NMT1 eQTL score** |  |  |  |  |  |
| Global MD | -0.0328 | 0.0075 | -4.3626 | 1.2941E-05 | 0.0257 |
| **CPNE1 eQTL score** |  |  |  |  |  |
| Global MD | -0.0366 | 0.0075 | -4.8650 | 1.1564E-06 | 0.0050 |
| **PLEKHM1 eQTL score** |  |  |  |  |  |
| Global MD | 0.0330 | 0.0075 | 4.3859 | 1.1631E-05 | 0.0250 |

**Table 6.** Significant associations between eQTL scores and tract categories (MD); FDR = false discovery rate; Effect size in table is ordered from smallest to largest for each score.

| Score, White Matter Tracts | Effect size | SD | t value | p value | p value, FDR corrected |
| --- | --- | --- | --- | --- | --- |
| **DCAKD eQTL score** |  |  |  |  |  |
| Thalamic radiations | 0.0327 | 0.0072 | 4.5625 | 5.09715E-06 | 0.0132 |
| Association fibres | 0.0381 | 0.0077 | 4.9643 | 6.97256E-07 | 0.0045 |
| **CPNE1 eQTL score** |  |  |  |  |  |
| Association fibres | -0.0368 | 0.0077 | -4.7868 | 1.7111E-06 | 0.0063 |
| **PLEKHM1 eQTL score** |  |  |  |  |  |
| Thalamic radiations | 0.0296 | 0.0072 | 4.1282 | 3.6762E-05 | 0.0423 |
| Association fibres | 0.0318 | 0.0077 | 4.1395 | 3.5002E-05 | 0.0423 |

**Table 7.** Significant associations between eQTL scores and individual white matter tracts (MD); FDR = false discovery rate; Effect size in table is ordered from smallest to largest for each score.

| Score, White Matter Tracts | Effect size | SD | t value | p value | p value, FDR corrected |
| --- | --- | --- | --- | --- | --- |
| **DCAKD eQTL score** |  |  |  |  |  |
| Acoustic radiation | 0.0295 | 0.0069 | 4.2470 | 2.17989E-05 | 0.0472 |
| Uncinate fasciculus | 0.0314 | 0.0068 | 4.6086 | 4.08933E-06 | 0.0162 |
| Cingulate gyrus | 0.0352 | 0.0073 | 4.7887 | 1.69554E-06 | 0.0085 |
| Inferior longitudinal fasciculus | 0.0377 | 0.0073 | 5.1766 | 2.28961E-07 | 0.0024 |
| Anterior thalamic radiations | 0.0403 | 0.0070 | 5.7964 | 6.91525E-09 | 0.0003 |
| Inferior fronto-occipital fasciculus | 0.0410 | 0.0075 | 5.4805 | 4.31258E-08 | 0.0008 |
| Superior longitudinal fasciculus | 0.0415 | 0.0076 | 5.4902 | 4.08256E-08 | 0.0008 |
| Forceps minor | 0.0480 | 0.0076 | 6.3085 | 2.89925E-10 | 2.76005E-05 |
| **SLC35A4 eQTL score** |  |  |  |  |  |
| Inferior longitudinal fasciculus | 0.0362 | 0.0073 | 4.9676 | 6.8552E-07 | 0.0044 |
| Forceps minor | 0.0432 | 0.0076 | 5.6773 | 1.3949E-08 | 0.0004 |
| **SEC14L4 eQTL score** |  |  |  |  |  |
| Cingulate gyrus | 0.0328 | 0.0073 | 4.4648 | 8.074E-06 | 0.0248 |
| Acoustic radiation | 0.0339 | 0.0069 | 4.8778 | 1.0844E-06 | 0.0060 |
| Forceps minor | 0.0348 | 0.0076 | 4.5604 | 5.1479E-06 | 0.0188 |
| **SRA1 eQTL score** |  |  |  |  |  |
| Forceps minor | 0.0353 | 0.0076 | 4.6349 | 3.6022E-06 | 0.0155 |
| **NMT1 eQTL score** |  |  |  |  |  |
| Inferior longitudinal fasciculus | -0.0311 | 0.0073 | -4.2695 | 1.9718E-05 | 0.0447 |
| Inferior fronto-occipital fasciculus | -0.0335 | 0.0075 | -4.4845 | 7.3652E-06 | 0.0234 |
| Anterior thalamic radiations | -0.0339 | 0.0070 | -4.8703 | 1.1263E-06 | 0.0060 |
| Superior longitudinal fasciculus | -0.0343 | 0.0076 | -4.5355 | 5.7939E-06 | 0.0204 |
| Forceps minor | -0.0392 | 0.0076 | -5.1537 | 2.5879E-07 | 0.0025 |
| **CPNE1 eQTL score** |  |  |  |  |  |
| Inferior longitudinal fasciculus | -0.0309 | 0.0073 | -4.2303 | 2.3485E-05 | 0.0497 |
| Superior longitudinal fasciculus | -0.0356 | 0.0076 | -4.7055 | 2.5555E-06 | 0.0116 |
| **PLEKHM1 eQTL score** |  |  |  |  |  |
| Forceps minor | 0.0334 | 0.0076 | 4.3876 | 1.154E-05 | 0.0297 |
| Superior longitudinal fasciculus | 0.0342 | 0.0076 | 4.5223 | 6.1651E-06 | 0.0210 |
| Anterior thalamic radiations | 0.0356 | 0.0070 | 5.1101 | 3.2604E-07 | 0.0028 |
| **UBE3C eQTL score** |  |  |  |  |  |
| Forceps minor | 0.0331 | 0.0076 | 4.3465 | 1.3925E-05 | 0.0349 |
| Inferior fronto-occipital fasciculus | 0.0332 | 0.0075 | 4.4413 | 9.01E-06 | 0.0268 |

**Figure 2a and 2b.**
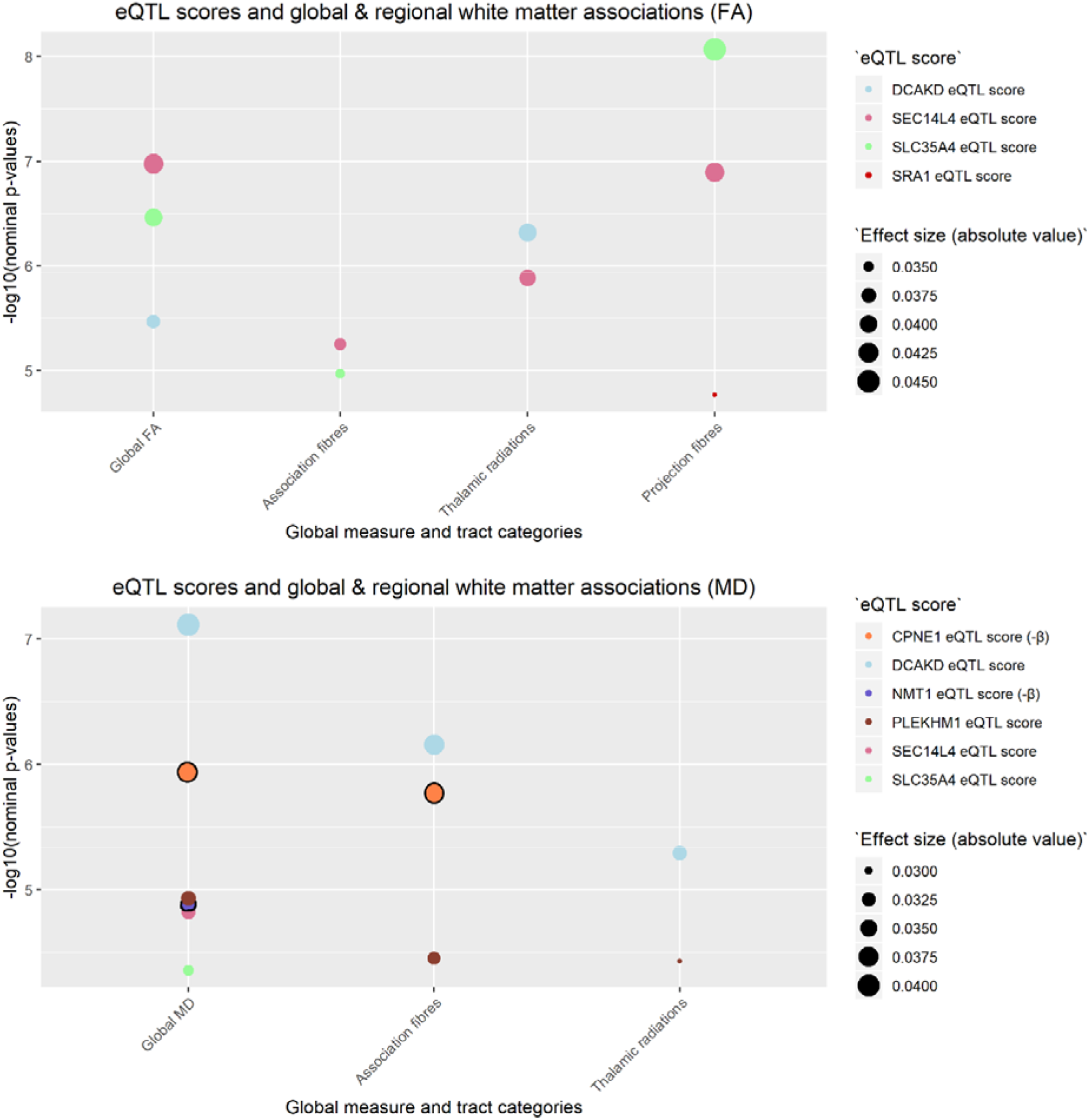
Indicates nominal p-values between each of the 8 scores (shown in legend entitled “eQTL score”) and global and tract category measures (noted on the x-axis; FA = fractional anisotropy (figure 2a); MD = mean diffusivity (figure 2b)). All values in the figure met FDR correction. Two of the scores with an additional line around the points (CPNE1 and NMT1) had an effect size in the opposite direction to all other scores (also indicated by -β for MD in figure legend). The colours of the plot points indicate the score to which they belong. Magnitude of effect is shown in the legend entitled “Effect size (absolute values)”.

**Figure 3a and 3b.**
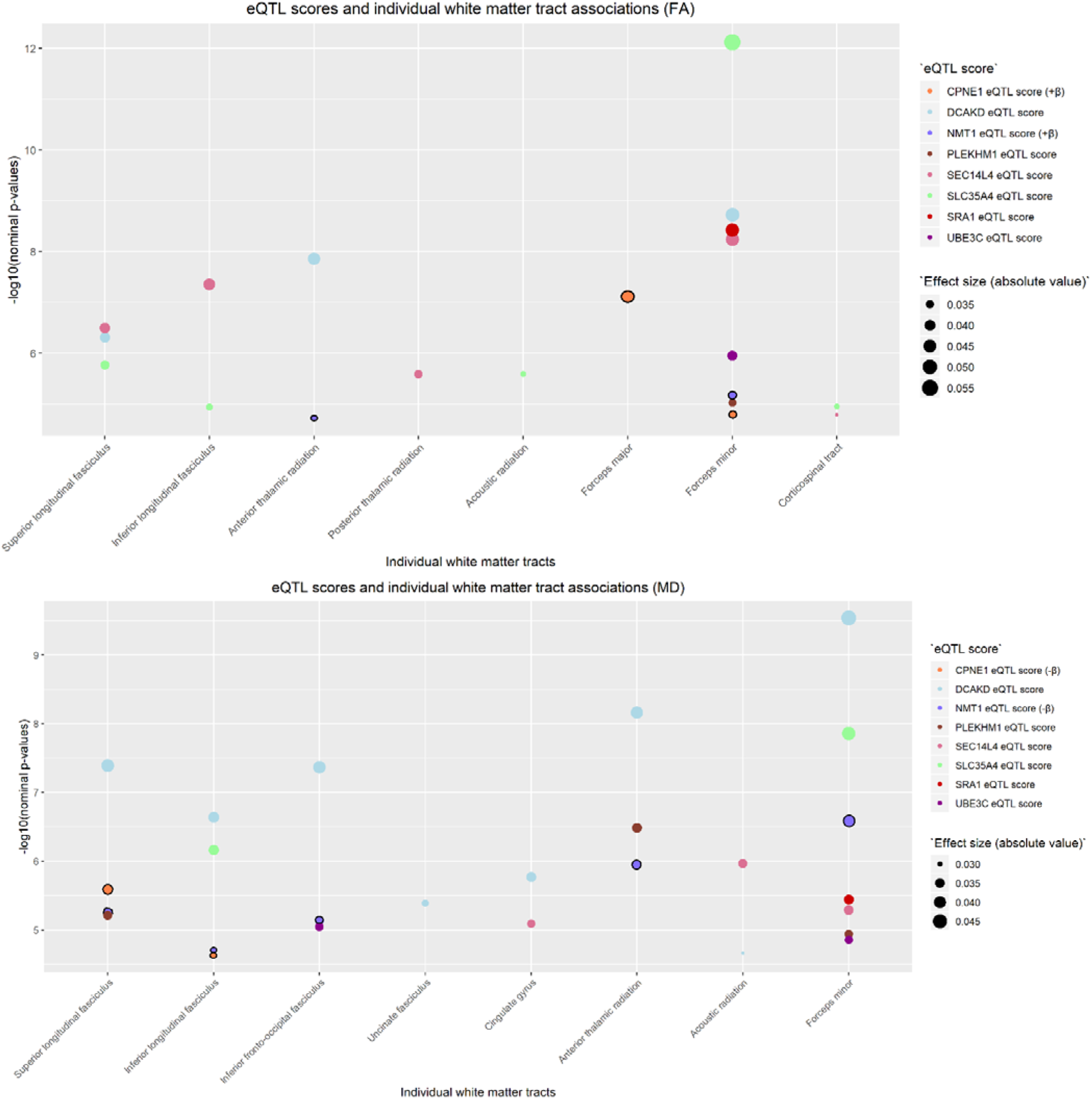
Indicates nominal p-values between each of the 8 scores (shown in legend entitled “eQTL score”) and individual white matter tracts (noted on the x-axis; FA = fractional anisotropy (figure 3a); MD = mean diffusivity (figure 3b)). All values in the figure met FDR correction. Two of the scores with an additional line around the points (CPNE1 and NMT1) had an effect size in the opposite direction to all other scores (+β and −β for FA and MD, respectively in figure legend). The colours of the plot points indicate the score to which they belong. Magnitude of effect is shown in the legend entitled “Effect size (absolute values)”.

### Genome-wide associations between score SNPs and white matter tracts

Using a previously published GWAS of imaging traits (27), we next sought to observe the association between the SNPs comprising each of the 8 scores (N_total_ = 53; SNP list can be found in section 7 of supplementary materials) with those found previously for the white matter tracts of interest (i.e. the tracts which showed post-FDR significant associations). This SNP look-up was performed in order to observe whether our analysis of eQTL scores, comprising SNPs which together regulate the expression of a single gene, yielded any novel associations with white matter tracts which were not previously found in conventional GWAS.

We used the Brain Imaging Genetics (BIG) database (http://big.stats.ox.ac.uk/) to extract the effect size and p-value of each SNP of interest as associated with the white matter tracts of interest, as provided in Elliott et al. (27). As GWAS for global and tract category measures were not performed in the original study (27), we performed these GWAS as part of the current project (i.e. GWAS for global measures, association fibres, thalamic radiations and projection fibres). Our GWAS parameters and quality check procedures are described in more detail in supplementary materials, section 4. P-values and effect size of each SNP for each individual white matter tract of interest (left and right hemispheres separately from Elliott et al. (27)), as well as for global and tract categories (run locally), are also contained in supplementary materials, section 6. Briefly, only one SNP across three eQTL scores was previously found to reach genome-wide significance with forceps minor (FA) (27), projection fibres (FA) and global FA (GWAS run locally): rs2237077.

## Discussion

In this study, we utilised a novel approach to investigate whether eQTL scores corresponding to the expression of specific genes in whole blood were significantly and specifically associated with white matter tracts in N > 14,000 individuals. We found significant associations in white matter microstructure as measured by both FA and MD for a number of scores (FA_N scores_ = 25; MD_N scores_ = 24). Of these, 8 scores were found to be significantly associated with various white matter tracts as measured by both FA and MD. In particular, the largest effect was seen for the association between forceps minor (FA) and the eQTL score for *SLC35A4*, and across several tracts measured by MD for the eQTL score for *DCAKD*. Although these eQTL were derived from whole blood, there is evidence of expression in the brain for some of the genes, outlined in further detail below. These findings also provided novel information not previously found by conventional genome-wide association studies.

All 8 scores were associated with white matter microstructural integrity of the forceps minor as measured by FA (7 of which were also associated with MD values). The forceps minor forms the anterior part of the corpus callosum, connecting homologous regions of the prefrontal cortex between hemispheres. It is postulated to be involved in numerous cognitive and behavioural skills, such as decision making, social behaviour, and language (28). This connection therefore implicates forceps minor in a wide range of cognitive skills, and damage to the tract has been associated with neuropsychiatric and neurological disorders, such as multiple sclerosis and depression (29; 30).

### Global and individual tract findings – largest associations

The two genes with the largest associations were *DCAKD*, globally and across numerous tracts as measured by higher MD, and *SCL35A4* across tracts measured by lower FA, with a peak in projection fibres, localised to forceps minor. *DCAKD* is a protein coding gene which is ubiquitously expressed in brain, among other tissues (31). Previous evidence using mouse models indicates expression of this gene has a putative role in neurodevelopment (32), and is associated with a number of psychiatric and neurological disorders, including schizophrenia, autism spectrum disorder, and Parkinson’s disease (31; 33; 34). Evidence for involvement in autism spectrum disorder comes from Butler et al. (2015) (34), who compiled a list of clinically relevant genes for the disorder, with *DCAKD* among the participating susceptibility genes. Expression of *DCAKD* was also found to be implicated in Parkinson’s disease (31), a disorder previously associated with lower white matter integrity in tracts within the temporal, parietal and occipital lobes (35).

*SLC35A4* belongs to the *SLC35* family, members of which act as transporters of nucleotide sugars, and is known to be expressed in brain (36). There is limited knowledge about its specific function, although a recent review investigating the subcellular localization and topology of *SLC35A4* demonstrated that it localizes mainly to the Golgi apparatus (36).

### Disease-linked genes - lower FA & higher MD (decreased white matter integrity)

Four genes (*SRA1, UBE3C, SEC14L4, PLEKHM1*) were associated with lower FA within several individual tracts pertaining to projection and association fibres, as well as with higher global MD. *SRA1* encodes both non-coding and protein-coding RNAs, is implicated in the regulation of numerous nuclear receptor activities, such as metabolism and chromatin organization, and is known to be expressed in the brain. Kotan et al. (2016) (37) posited that *SRA1* plays a role in the initiation of puberty in humans by finding that inactivating *SRA1* variants were associated with idiopathic hypogonadotropic hypogonadism (IHH) in three independent families. IHH is a rare genetic disorder caused by the inability of the hypothalamus to secrete gonadotropin-releasing hormones (GnRH) or by the inability of GnRH to act on pituitary gonadotropes (38). These previous results might link the association of *SRA1* with projection fibres, which connect the cerebral cortex to the spinal cord and brainstem, as well as to other centres of the brain (e.g. thalamus).

*UBE3C* contains ubiquitin-protein ligase (E3), an enzyme which accepts ubiquitin from E2 before transferring it to the target lysine; ubiquitin targets proteins for degradation via the proteasome. *UBE3C* is expressed in numerous tissues, including the brain, and has been previously associated with some neuropsychiatric-related phenotypes. For instance, Garriock et al. (2010) (39) performed a GWAS to determine the association between genetic variation and Citalopram response. Although not genome-wide significant, their top finding was a SNP in proximity to *UBE3C* and was found to be associated with antidepressant response and MDD remission (rs6966038, p = 4.65e-07 and p = 3.63E-07, respectively) (39). Moreover, Filatova et al. (2014) (40) studied the expression of genes within the ubiquitin-proteasome protein degradation system, which is implicated in Parkinson’s disease, in mice with MPTP-induced pre-symptomatic and early symptomatic stages of Parkinson’s disease. They found decreased expression in the striatum and the substantia nigra of mice, which may lead to a decrease in performance of the system. This may in turn lead to accumulation of abnormal and toxic proteins which guide neuronal cell death (40).

The specific function of *SEC14L4* has not yet been determined, although the protein encoded by it is similar to a protein encoded by the *SEC14* gene in saccharomyces cerevisiae, which is essential to the biogenesis of Golgi-derived transport vesicles. Curwin and McMaster (2008) (41) found that mutations in several *SEC14* domain-containing proteins in humans may be implicated in neurodegeneration, although it is not clear what the role of *SEC14L4* is within this context. Lastly, *PLEKHM1* is important in bone resorption, may be involved in vesicular transport in the osteoclast, and is weakly expressed in the brain. Although mutations in this gene have been associated with numerous phenotypes (42; 43), none were neuropsychiatric-related.

### Development-linked genes - higher FA & lower MD (increased white matter integrity)

For two of the eight genes (*CPNE1, NMT1*) we found higher FA and lower MD, indicating increased white matter integrity, associated with increased expression level as quantified by the corresponding eQTL.

*CPNE1*, which is thought to regulate molecular events at the cell membrane and cytoplasm, has previously been found to mediate several neuronal differentiation processes by interacting with intracellular signalling molecules. *CPNE1* has also been found to be highly expressed during brain development, indicating that it might be implicated in earlier developmental stages of neuronal function (44). Furthermore, C2 domains of *CPNE1*, calcium-dependent phospholipid-binding motives, have been shown to be implicated in neurite outgrowth of hippocampal progenitor HiB5 cells, which are hippocampal cell lines derived from the hippocampal analgen of E16 rat (45; 46). We provide evidence that *CPNE1* expression is associated with two tracts within projection fibres (FA) and with regional association fibres (MD), which link the cortex to lower brain areas. In mouse and human models, these findings may be of use when investigating neurite outgrowth from the hippocampus, which is part of the limbic system, an area located beneath the cortex.

*NMT1* (N-myristoyltransferase) catalyzes the transfer of myristate (a rare 14-carbon saturated fatty acid) from CoA to proteins, and is expressed in numerous tissues, including ubiquitously in the brain. It has been found that *NMT1* is required for early mouse development, mainly due to its role in early embryogenesis (47). Expression of this gene has also been implicated in human brain tumours (48) and tumour cell proliferation (49). In our study, we found *NMT1* to be associated with tracts within thalamic radiations and projection fibres (FA) and global MD.

### General Discussion

In our study, we employed a novel strategy of investigating a direct association between eQTL scores and white matter tracts to uncover a relationship between specific regulatory variants and brain connectivity. Together, our findings indicate that increases in expression of these genes may be implicated in several processes which may directly or indirectly alter white matter microstructure, each with localised, pronounced effects in specific tracts. Further, while some of the significant associations had connections with other brain-related traits, such as neurite outgrowth or psychiatric and neurological disorders, others did not. Interestingly, decreased white matter microstructure integrity, as marked by lower FA and higher MD, was associated with eQTL scores which regulate expression of genes implicated in neuropsychiatric and neurological disorders. Conversely, increased white matter integrity, as marked by higher FA and lower MD, was associated with *CPNE1* and *NMT1*, which are important in developmental processes such as neurite outgrowth. In addition, encouragingly, regions of the corpus callosum (i.e. the forceps minor), the largest and arguably most reliably measured white matter tract in the brain, was demonstrated to be associated with all 8 scores for FA, and 7 for MD. These findings together suggest that utilising this approach to associate eQTL scores with white matter microstructure may add to previous research which found associations between genes and these brain-related traits and disorders. These genes or eQTL for them might indirectly implicate brain connectivity through other processes in which they participate.

The current study has several strengths and some potential limitations. First, to our knowledge, this study is the first one to compute eQTL scores for specific gene transcripts and attempt to associate them with white matter tracts. Moreover, our analysis consisted of a population-based sample of N > 14,000 individuals recruited to the UKB, large enough to make our findings robust and generalizable to other samples within the same age range, background and ethnicity. Lastly, our findings revealed novel associations which were not previously found in GWAS (27; GWAS of g measures run locally), indicating a potential to use such scores for further discovery analyses.

However, a potential limitation in this study is calculation of scores for data taken from whole blood. Although we note previous evidence indicates that whole blood can be used as a proxy for brain eQTL, important for study of *in vivo* brain traits (14).

In summary, our results suggest that expression of the genes discussed above alter white matter microstructure and could facilitate the manifestation of numerous brain-related traits. Uncovering specific markers leading to the formation, maintenance and pathology of white matter could enable downstream analyses to elucidate links between genetics and neuroimaging in neurological and psychiatric disorders, as well as other brain-related traits.

