## Supplementary materials for "Expression quantitative trait loci-derived scores and white matter microstructure in UK Biobank: a novel approach to integrating genetics and neuroimaging"

1. **Results for scores associated with FA (N = 17; table 1) and MD (N = 16; table 2) white matter tracts separately**

| **Score, white matter tract** | **Effect size** | **SD** | **t value** | **p value** | **p value, FDR corrected** |
| --- | --- | --- | --- | --- | --- |
| ATG10_eQTL_cis, global FA | 0.0341 | 0.0079 | 4.3106 | 1.64E-05 | 0.0273 |
| SF3A1_eQTL_cis, global FA | -0.0327 | 0.0079 | -4.1305 | 3.64E-05 | 0.0495 |
| SMARCAL1_eQTL_cis, global FA | 0.0374 | 0.0079 | 4.7354 | 2.21E-06 | 0.0071 |
| SF3A1_eQTL_cis, association fibres | -0.0334 | 0.0079 | -4.2386 | 2.26E-05 | 0.0344 |
| SMARCAL1_eQTL_cis, association fibres | 0.0326 | 0.0079 | 4.1375 | 3.53E-05 | 0.0495 |
| ATG10_eQTL_cis, thalamic radiations | 0.0373 | 0.0080 | 4.6587 | 3.21E-06 | 0.0088 |
| PPP4R3A_eQTL_cis, thalamic radiations | 0.0357 | 0.0080 | 4.4572 | 8.36E-06 | 0.0166 |
| SMARCAL1_eQTL_cis, thalamic radiations | 0.0394 | 0.0080 | 4.9292 | 8.35E-07 | 0.0036 |
| CD14_eQTL_cis, projection fibres | -0.0360 | 0.0079 | -4.5691 | 4.94E-06 | 0.0116 |
| COG7_eQTL_cis, anterior thalamic radiation | -0.0333 | 0.0076 | -4.4005 | 1.09E-05 | 0.0337 |
| SMARCAL1_eQTL_cis, anterior thalamic radiation | 0.0394 | 0.0076 | 5.2164 | 1.85E-07 | 0.0018 |
| LINC01605_eQTL_trans, cingulate gyrus | -0.0337 | 0.0071 | -4.7560 | 1.99E-06 | 0.0114 |
| ANXA1_eQTL_cis, corticospinal tract | -0.0320 | 0.0074 | -4.3218 | 1.56E-05 | 0.0416 |
| ZSCAN26_eQTL_cis, forceps major | -0.0397 | 0.0081 | -4.9048 | 9.45E-07 | 0.0070 |
| ATG10_eQTL_cis, forceps minor | 0.0360 | 0.0078 | 4.5986 | 4.29E-06 | 0.0189 |
| CD14_eQTL_cis, forceps minor | 0.0456 | 0.0078 | 5.8210 | 6E-09 | 0.0001 |
| SHTN1 / KIAA1598_eQTL_cis, forceps minor | 0.0376 | 0.0078 | 4.8050 | 1.56E-06 | 0.0101 |
| ZNF282_eQTL_cis, forceps minor | -0.0346 | 0.0078 | -4.4224 | 9.83E-06 | 0.0337 |
| ENO4_eQTL_cis, forceps minor | 0.0354 | 0.0078 | 4.5197 | 6.24E-06 | 0.0252 |
| COG7_eQTL_cis, forceps minor | -0.0338 | 0.0078 | -4.3127 | 1.62E-05 | 0.0416 |
| SMARCAL1_eQTL_cis, forceps minor | 0.0361 | 0.0078 | 4.6056 | 4.15E-06 | 0.0189 |
| ASRGL1_eQTL_cis, inferior fronto-occipital fasciculus | 0.0329 | 0.0077 | 4.2950 | 1.76E-05 | 0.0426 |
| ATG10_eQTL_cis, inferior fronto-occipital fasciculus | 0.0355 | 0.0077 | 4.6291 | 3.7E-06 | 0.0179 |
| TMEM184B_eQTL_cis, inferior fronto-occipital fasciculus | 0.0337 | 0.0077 | 4.3935 | 1.12E-05 | 0.0337 |
| SMARCAL1_eQTL_cis, inferior longitudinal fasciculus | 0.0349 | 0.0076 | 4.5704 | 4.91E-06 | 0.0207 |
| ATG10_eQTL_cis, posterior thalamic radiation | 0.0325 | 0.0075 | 4.3416 | 1.42E-05 | 0.0406 |
| ZBTB7B_eQTL_cis, superior longitudinal fasciculus | -0.0329 | 0.0077 | -4.2946 | 1.76E-05 | 0.0426 |
| GPT_eQTL_cis, superior longitudinal fasciculus | 0.0339 | 0.0077 | 4.4153 | 1.02E-05 | 0.0337 |
| SMARCAL1_eQTL_cis, superior longitudinal fasciculus | 0.0401 | 0.0077 | 5.2356 | 1.67E-07 | 0.0018 |
| GPT_eQTL_cis, superior thalamic radiation | 0.0337 | 0.0079 | 4.2827 | 1.86E-05 | 0.0429 |
| AP2S1_eQTL_cis, superior thalamic radiation | 0.0348 | 0.0079 | 4.4164 | 1.01E-05 | 0.0337 |

**Table 1.** eQTL scores associated only with white matter tracts as measured by FA.

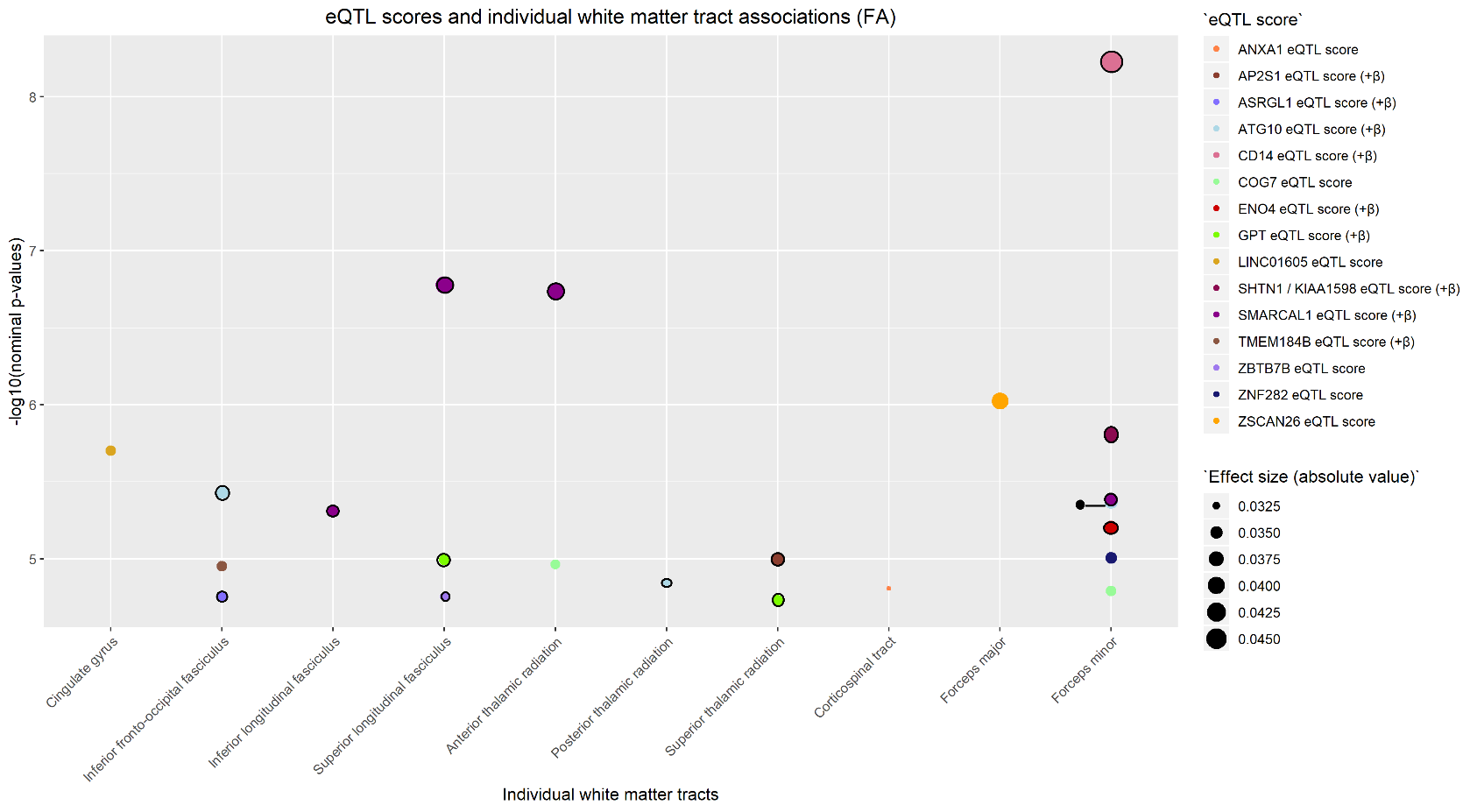

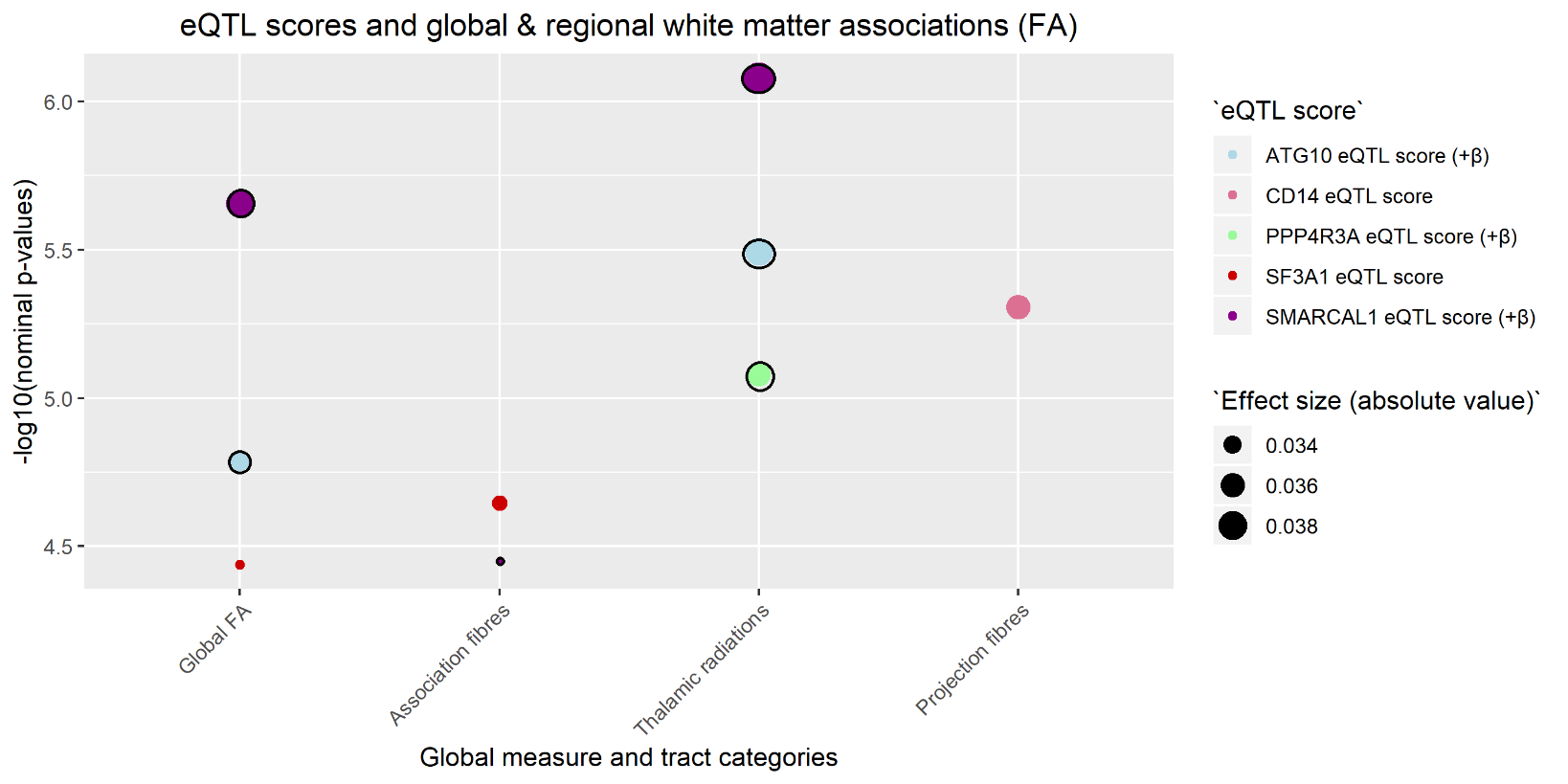

**Figure 1.** eQTL scores associated only with white matter tracts as measured by FA (fractional anisotropy). Indicates nominal p-values between each of the scores (shown in legend entitled “eQTL score”) and global and tract category measures (noted on the x-axis). All values in the figure met FDR correction. Some of the scores with an additional line around the points had an effect size in the opposite direction to all other scores (also indicated by +β for FA in figure legend). The colours of the plot points indicate the score to which they belong. Magnitude of effect is shown in the legend entitled “Effect size (absolute values)”.

| **Score, white matter tract** | **Effect size** | **SD** | **t value** | **p value** | **p value, FDR corrected** |
| --- | --- | --- | --- | --- | --- |
| APOA1BP / NAXE_eQTL_cis, global MD | 0.0311 | 0.0075 | 4.1331 | 3.6E-05 | 0.0423 |
| BTN3A2_eQTL_cis, global MD | 0.0308 | 0.0075 | 4.0853 | 4.42E-05 | 0.0423 |
| UMPS_eQTL_cis, global MD | -0.0319 | 0.0075 | -4.2381 | 2.27E-05 | 0.0366 |
| CSF3R_eQTL_cis, global MD | 0.0345 | 0.0075 | 4.5704 | 4.91E-06 | 0.0132 |
| TMEM154_eQTL_cis, global MD | -0.0400 | 0.0076 | -5.2970 | 1.19E-07 | 0.0015 |
| APOA1BP / NAXE_eQTL_cis, association fibres | 0.0326 | 0.0077 | 4.2419 | 2.23E-05 | 0.0366 |
| BTN3A2_eQTL_cis, association fibres | 0.0314 | 0.0077 | 4.0888 | 4.36E-05 | 0.0423 |
| SAMM50_eQTL_cis, association fibres | -0.0311 | 0.0077 | -4.0501 | 5.15E-05 | 0.0475 |
| UMPS_eQTL_cis, association fibres | -0.0355 | 0.0077 | -4.6219 | 3.84E-06 | 0.0124 |
| CSF3R_eQTL_cis, association fibres | 0.0377 | 0.0077 | 4.9069 | 9.36E-07 | 0.0048 |
| TMEM154_eQTL_cis, association fibres | -0.0402 | 0.0077 | -5.2158 | 1.86E-07 | 0.0016 |
| HLA-C_eQTL_cis, association fibres | -0.0342 | 0.0077 | -4.4402 | 9.05E-06 | 0.0213 |
| MED15_eQTL_cis, thalamic radiations | 0.0297 | 0.0072 | 4.1354 | 3.56E-05 | 0.0423 |
| KANSL1_eQTL_cis, thalamic radiations | -0.0302 | 0.0072 | -4.2041 | 2.64E-05 | 0.0401 |
| IL18RAP_eQTL_cis, projection fibres | 0.0324 | 0.0078 | 4.1621 | 3.17E-05 | 0.0423 |
| C6orf106_eQTL_cis, projection fibres | 0.0318 | 0.0078 | 4.0863 | 4.41E-05 | 0.0423 |
| RABEPK_eQTL_cis, acoustic radiation | 0.0307 | 0.0069 | 4.4197 | 9.95E-06 | 0.0287 |
| CFDP1_eQTL_cis, anterior thalamic radiation | 0.0298 | 0.0070 | 4.2749 | 1.92E-05 | 0.0447 |
| PTPN13_eQTL_cis, anterior thalamic radiation | -0.0347 | 0.0070 | -4.9756 | 6.58E-07 | 0.0044 |
| KANSL1_eQTL_cis, anterior thalamic radiation | -0.0369 | 0.0070 | -5.3005 | 1.17E-07 | 0.0016 |
| TMEM154_eQTL_cis, anterior thalamic radiation | -0.0303 | 0.0070 | -4.3269 | 1.52E-05 | 0.0372 |
| UMPS_eQTL_cis, cingulate gyrus | -0.0323 | 0.0073 | -4.3999 | 1.09E-05 | 0.0297 |
| PLEC_eQTL_cis, forceps minor | 0.0335 | 0.0076 | 4.3887 | 1.15E-05 | 0.0297 |
| TMEM154_eQTL_cis, forceps minor | -0.0384 | 0.0076 | -5.0145 | 5.38E-07 | 0.0043 |
| TMEM154_eQTL_cis, inferior fronto-occipital fasciculus | -0.0330 | 0.0075 | -4.3971 | 1.1E-05 | 0.0297 |
| TMEM154_eQTL_cis, inferior longitudinal fasciculus | -0.0337 | 0.0073 | -4.5974 | 4.32E-06 | 0.0164 |
| SAMM50_eQTL_cis, parahippocampal part of cingulum | -0.0303 | 0.0071 | -4.2755 | 1.92E-05 | 0.0447 |
| BTN3A2_eQTL_cis, superior longitudinal fasciculus | 0.0350 | 0.0076 | 4.6271 | 3.74E-06 | 0.0155 |
| UMPS_eQTL_cis, superior longitudinal fasciculus | -0.0413 | 0.0076 | -5.4562 | 4.94E-08 | 0.0008 |
| TMEM154_eQTL_cis, superior longitudinal fasciculus | -0.0377 | 0.0076 | -4.9633 | 7.01E-07 | 0.0044 |
| PTPN13_eQTL_trans, superior thalamic radiation | -0.0336 | 0.0068 | -4.9424 | 7.8E-07 | 0.0046 |
| TMEM154_eQTL_cis, superior thalamic radiation | -0.0323 | 0.0068 | -4.7333 | 2.23E-06 | 0.0106 |
| TMEM154_eQTL_cis, uncinate fasciculus | -0.0308 | 0.0068 | -4.5058 | 6.66E-06 | 0.0219 |

**Table 2.** eQTL scores associated only with white matter tracts as measured by MD.

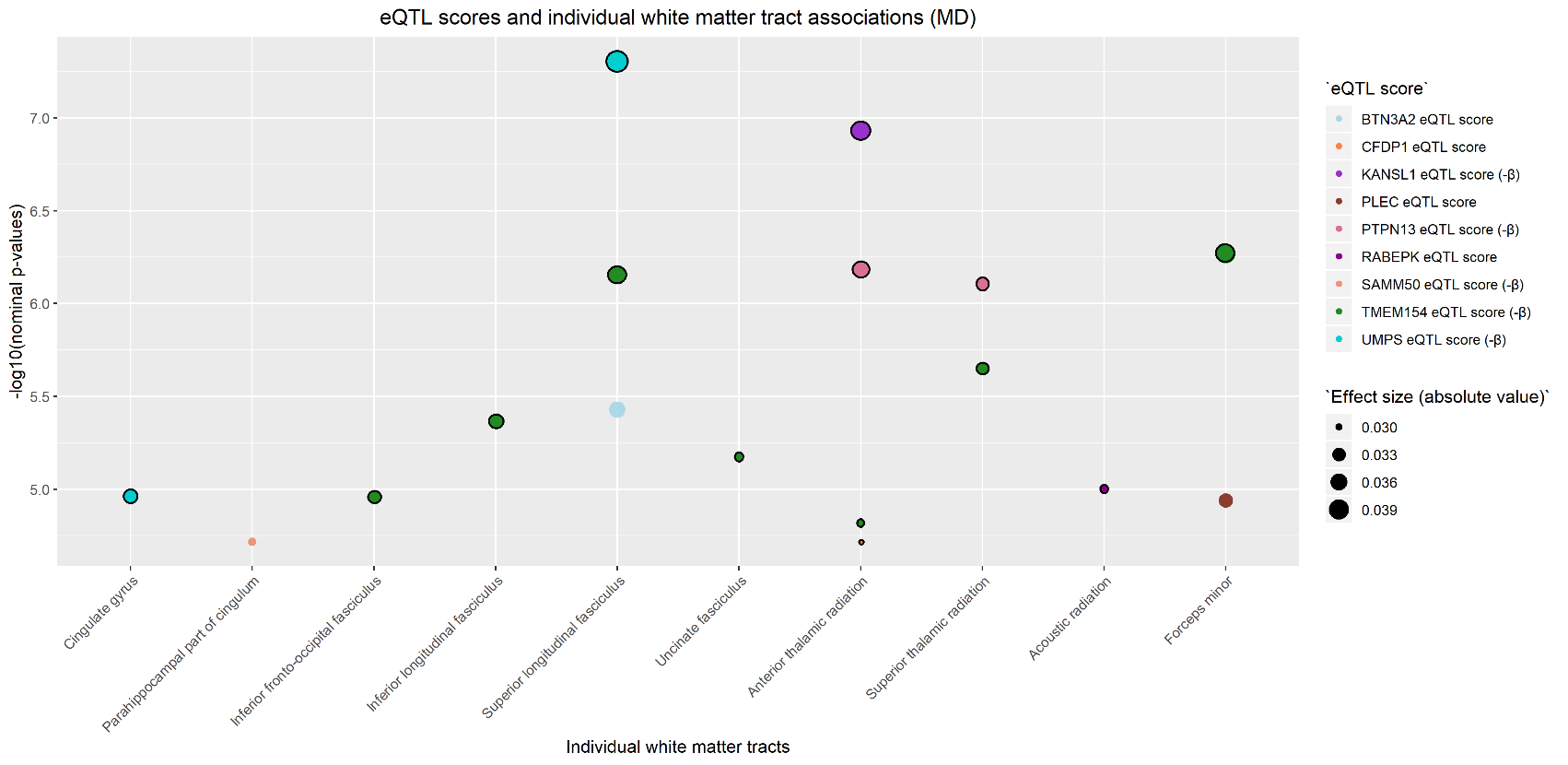

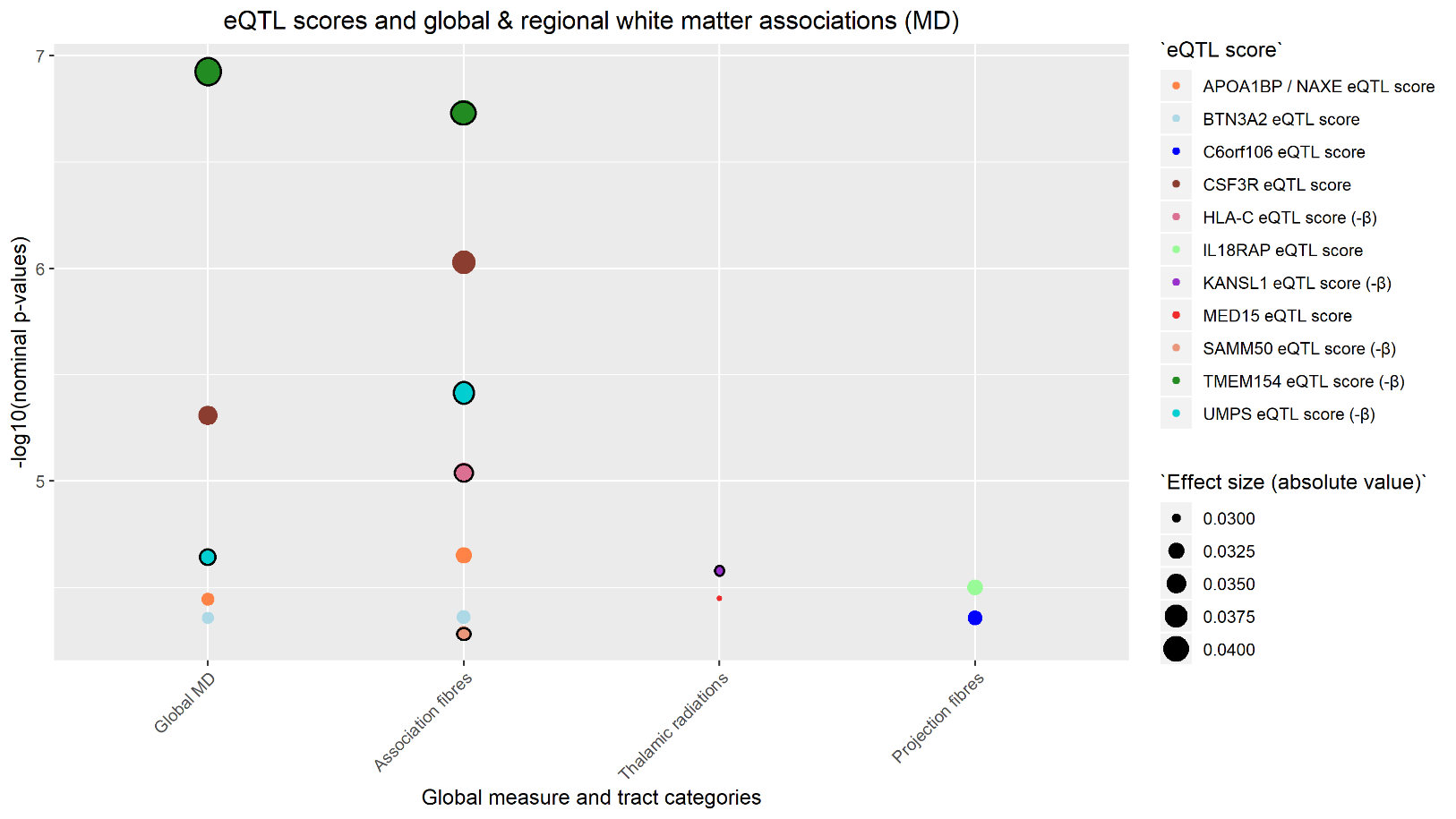

**Figure 2.** eQTL scores associated only with white matter tracts as measured by MD (mean diffusivity). Indicates nominal p-values between each of the scores (shown in legend entitled “eQTL score”) and global and tract category measures (noted on the x-axis). All values in the figure met FDR correction. Some of the scores with an additional line around the points had an effect size in the opposite direction to all other scores (also indicated by -β for MD in figure legend). The colours of the plot points indicate the score to which they belong. Magnitude of effect is shown in the legend entitled “Effect size (absolute value)”.

| 2. **Brief gene look-up for genes whose expression was associated with FA (N = 17; table 1) and MD (N = 16; table 2) separately** | | | | |
| --- | --- | --- | --- | --- |
| **Score name & eQTL type** | **N SNPs in score** | **Regulated gene** | **Study from which score is calculated** | **Gene function** |
| ATG10_eQTL_cis | 7 | ATG10 | Gusev et al. | E2-like enzyme involved in 2 ubiquitin-like modifications essential for autophagosome formation; expressed in brain (1) |
| SF3A1_eQTL_cis | 23 | SF3A1 | Westra et al. | Expressed in brain; gene encodes a subunit of the splicing factor 3a protein complex (2) |
| SMARCAL1_eQTL_cis | 1 | SMARCAL1 | Westra et al. | Protein encoded by this gene is a member of SWI/SNP family of proteins; members have helicase and ATPase activities and are thought to regulate transcription of certain genes by altering chromatin structure around those genes; expressed in brain; associated with Schimke immunoosseous dysplasia (3) |
| PPP4R3A_eQTL_cis | 1 | PPP4R3A | Westra et al, | Expressed in brain; may be involved in Alzheimer’s disease risk (4) |
| CD14_eQTL_cis | 18 | CD14 | Gusev et al. | Protein encoded by this gene is a surface antigen that is preferentially expressed on monocytes/macrophages; it cooperates with other proteins to mediate the innate immune response to bacterial lipopolysaccharide; expressed in brain (5) |
| COG7_eQTL_cis | 1 | COG7 | Westra et al. | [Protein encoded by this gene resides in the golgi and is part of 8 subunits of the conserved oligomeric Golgi (COG) complex; expressed in brain; mutations in gene associated with microcephaly, adducted thumbs, growth retardation, VSD and episodes of hyperthermia (6; 7)](Protein%20encoded%20by%20this%20gene%20resides%20in%20the%20golgi%20and%20is%20part%20of%208%20subunits%20of%20the%20conserved%20oligomeric%20Golgi%20(COG)%20complex;%20expressed%20in%20brain;%20) |
| LINC01605_eQTL_trans | 1 | LINC01605 | Westra et al. | [RNA gene; affiliated with non-coding RNA class; expression of gene associated with bladder cancer](http://www.bioscirep.org/content/early/2018/07/27/BSR20180562) (8) |
| ANXA4_eQTL_cis | 5 | ANXA4 | Gusev et al. | Little expression in brain; gene belongs to annexin family of calcium dependent phospholipid binding proteins (9) |
| ZSCAN26_eQTL_cis | 32 | ZSCAN26 | Westra et al. | Expressed in brain; protein coding gene |
| SHTN1 / KIAA1598_eQTL_cis | 5 | SHTN1 / KIAA1598 | Gusev et al. | [Expressed in brain; involved in generation of internal asymmetric signals required for neuronal polarization and neurite outgrowth; mediated netrin-1-induced F-actin substrate coupling or clutch engagement within axon growth cone through activation of several genes & pathways](https://www.ncbi.nlm.nih.gov/pubmed/18519736/) (10) |
| ZNF282_eQTL_cis | 7 | ZNF282 | Gusev et al. | Expressed in brain; diseases associated with gene: T-cell leukemia (11) |
| ENO4_eQTL_cis | 7 | ENO4 | Westra et al. | Expressed in brain |
| ASRGL1_eQTL_cis | 5 | ASRGL1 | Gusev et al. | Expressed in brain; may be involved in production of L-aspartate, which can act as an excitatory neurotransmitter in some brain regions; may be implicated in endometrioid endometrial carcinoma (12) |
| TMEM184B_eQTL_cis | 12 | TMEM184B | Gusev et al. | Expressed in brain; may be implicated in axon degeneration (13) |
| ZBTB7B_eQTL_cis | 8 | ZBTB7B | Gusev et al. | Expressed in brain; gene encodes a zinc finger-containing transcription factor that acts as a key regulator of lineage commitment of immature T-cell precursors (14) |
| GPT_eQTL_cis | 5 | GPT | Westra et al. | Little expression in brain |
| AP2S1_eQTL_cis | 5 | AP2S1 | Westra et al. | One of 2 major clathrin-associated adaptor complexes, AP-2 is a heterotetramer which is associated with the plasma membrane; complex is composed of 2 large chains, 1 medium chain and 1 small chain, and the gene encodes the small chain; expressed in brain (15) |

**Table 1**. Information regarding eQTL scores with significant associations FA-measured tracts. 1 score (LINC01605_eQTL_trans) is trans, while all others are cis.

| **Score name & eQTL type** | **N SNPs in score** | **Regulated gene** | **Study from which score is calculated** | **Gene function** |
| --- | --- | --- | --- | --- |
| APOA1BP / NAXE_eQTL_cis | 10 | APOA1BP / NAXE | Gusev et al. | [Expressed in brain; diseases associated with gene: encephalopathy; brain edema](https://www.ncbi.nlm.nih.gov/pubmed/27616477/) (16) |
| BTN3A2_eQTL_cis | 42 | BTN3A2 | Gusev et al. | May be involved in adaptive immune system response; may be involved in risk for gastric cancer (17) |
| UMPS_eQTL_cis | 5 | UMPS | Westra et al. | Encoded protein is a bifunctional enzyme that catalyzes the final 2 steps of the de novo pyrimidine biosynthetic pathway (18) |
| CSF3R_eQTL_cis | 5 | CSF3R | Westra et al. | Mutations in this gene are a cause of Kostmann syndrome / congenital neutropenia; not expressed in brain (19) |
| TMEM154_eQTL_cis | 20 | TMEM154 | Westra et al. | Very little expression in brain |
| SAMM50_eQTL_cis | 15 | SAMM50 | Gusev et al. | Gene encodes a component of the Sorting and Assembly Machinery of the mitochondrial outer membrane (20) |
| HLA-C_eQTL_cis | 38 | HLA-C | Westra et al. | Expressed in nearly all cells |
| MED15_eQTL_cis | 6 | MED15 | Westra et al. | Expressed in brain |
| KANSL1_eQTL_cis | 3 | KANSL1 | Westra et al. | [Gene encodes a nuclear protein that is a subunit of 2 protein complexes involved with histone acetylation](https://bmcmedgenet.biomedcentral.com/articles/10.1186/s12881-015-0211-0) (21) |
| IL18RAP_eQTL_cis | 12 | IL18RAP | Gusev et al. | Mutations in this gene have been associated with Crohn's disease; expressed in brain (22) |
| C6orf106_eQTL_cis | 5 | C6orf106 | Westra et al. | [Expressed in cortex](https://www.proteinatlas.org/ENSG00000196821-C6orf106/tissue) |
| RABEPK_eQTL_cis | 9 | RABEPK | Gusev et al. | Expressed in brain |
| CFDP1_eQTL_cis | 4 | CFDP1 | Gusev et al. | [Expressed in brain; may be implicated in coronary artery disease risk](https://www.malacards.org/card/acute_stress_disorder) (23) |
| PTPN13_eQTL_trans | 1 | PTPN13 | Westra et al. | [Protein encoded by this gene is a member of the PTP family, which are signalling molecules that regulate cellular processes (e.g. cell growth, differentiation, mitotic cell cycle, oncogenic transformation); disease associated with this gene: tropical spastic paraparesis, a disease of the nervous system affecting people living near the equator; expressed in the brain](https://www.malacards.org/card/tropical_spastic_paraparesis) (24) |
| EVL_eQTL_cis | 1 | EVL | Westra et al. | Expressed in brain; actin-associated proteins involved in proccesses such as axon guidance and lamellipodial and filopodial dynamics in migrating cells; enhances actin nucleation and polymerization (25) |
| PLEC_eQTL_cis | 3 | PLEC | Westra et al. | Prominent member of a protein family of proteins which interlink different elements of the cytoskeleton; expressed in a wide range of cell types and tissues (including brain) (26) |

**Table 2**. Information regarding eQTL scores with significant associations MD-measured tracts. 1 score (PTPN13_eQTL_trans) is trans, while all others are cis.

3. **Results for 8 scores associated with both FA and MD (N = 8).**

| **White Matter Tracts** | **Effect size** | **SD** | **t value** | **p value** | **p value, FDR corrected** |
| --- | --- | --- | --- | --- | --- |
| **FA** |  |  |  |  |  |
| Global FA | -0.0367 | 0.0079 | -4.6474 | 3.39161E-06 | 0.0088 |
| Thalamic radiations | -0.0403 | 0.0080 | -5.0378 | 4.76577E-07 | 0.0025 |
| Anterior thalamic radiations | -0.0429 | 0.0076 | -5.6798 | 1.37465E-08 | 0.0002 |
| Forceps minor | -0.0471 | 0.0078 | -6.0115 | 1.88218E-09 | 0.0001 |
| Superior longitudinal fasciculus | -0.0386 | 0.0077 | -5.0327 | 4.89475E-07 | 0.0040 |
| **MD** |  |  |  |  |  |
| Global MD | 0.0404 | 0.0075 | 5.3762 | 7.72382E-08 | 0.0015 |
| Association fibres | 0.0381 | 0.0077 | 4.9643 | 6.97256E-07 | 0.0045 |
| Thalamic radiations | 0.0327 | 0.0072 | 4.5625 | 5.09715E-06 | 0.0132 |
| Acoustic radiation | 0.0295 | 0.0069 | 4.2470 | 2.17989E-05 | 0.0472 |
| Anterior thalamic radiations | 0.0403 | 0.0070 | 5.7964 | 6.91525E-09 | 0.0003 |
| Cingulate gyrus | 0.0352 | 0.0073 | 4.7887 | 1.69554E-06 | 0.0085 |
| Forceps minor | 0.0480 | 0.0076 | 6.3085 | 2.89925E-10 | 2.76005E-05 |
| Inferior fronto-occipital fasciculus | 0.0410 | 0.0075 | 5.4805 | 4.31258E-08 | 0.0008 |
| Inferior longitudinal fasciculus | 0.0377 | 0.0073 | 5.1766 | 2.28961E-07 | 0.0024 |
| Superior longitudinal fasciculus | 0.0415 | 0.0076 | 5.4902 | 4.08256E-08 | 0.0008 |
| Uncinate fasciculus | 0.0314 | 0.0068 | 4.6086 | 4.08933E-06 | 0.0162 |

**Table 1**. Significant associations between DCAKD_eQTL_cis and FA and MD-measured white matter tracts.

| **White Matter Tracts** | **Effect size** | **SD** | **t value** | **p value** | **p value, FDR corrected** |
| --- | --- | --- | --- | --- | --- |
| **FA** |  |  |  |  |  |
| Global FA | -0.0403 | 0.0079 | -5.0996 | 3.44595E-07 | 0.0022 |
| Association fibres | -0.0347 | 0.0079 | -4.4036 | 1.07241E-05 | 0.0198 |
| Projection fibres | 0.0453 | 0.0079 | 5.7612 | 8.51978E-09 | 0.0002 |
| Acoustic radiation | -0.0326 | 0.0069 | -4.7044 | 2.56987E-06 | 0.0133 |
| Corticospinal tract | -0.0326 | 0.0074 | -4.3945 | 1.11801E-05 | 0.0337 |
| Forceps minor | -0.0561 | 0.0078 | -7.1754 | 7.5595E-13 | 7.3217E-08 |
| Inferior longitudinal fasciculus | -0.0335 | 0.0076 | -4.3887 | 1.14829E-05 | 0.0337 |
| Superior longitudinal fasciculus | -0.0367 | 0.0077 | -4.7887 | 1.6956E-06 | 0.0103 |
| **MD** |  |  |  |  |  |
| Global MD | 0.0308 | 0.0075 | 4.0893 | 4.3502E-05 | 0.0423 |
| Forceps minor | 0.0432 | 0.0076 | 5.6773 | 1.3949E-08 | 0.0004 |
| Inferior longitudinal fasciculus | 0.0362 | 0.0073 | 4.9676 | 6.8552E-07 | 0.0044 |

**Table 2.** Significant associations between SLC35A4_eQTL_cis and FA and MD-measured white matter tracts.

| **White Matter Tracts** | **Effect size** | **SD** | **t value** | **p value** | **p value, FDR corrected** |
| --- | --- | --- | --- | --- | --- |
| **FA** |  |  |  |  |  |
| Global FA | -0.0420 | 0.0079 | -5.3199 | 1.0538E-07 | 0.0011 |
| Association fibres | -0.0358 | 0.0079 | -4.5425 | 5.6047E-06 | 0.0121 |
| Thalamic radiations | -0.0388 | 0.0080 | -4.8429 | 1.2928E-06 | 0.0048 |
| Projection fibres | 0.0416 | 0.0079 | 5.2850 | 1.275E-07 | 0.0011 |
| Corticospinal tract | -0.0320 | 0.0074 | -4.3116 | 1.6311E-05 | 0.0416 |
| Forceps minor | -0.0456 | 0.0078 | -5.8270 | 5.763E-09 | 0.0001 |
| Inferior longitudinal fasciculus | -0.0419 | 0.0076 | -5.4773 | 4.3905E-08 | 0.0006 |
| Posterior thalamic radiation | -0.0352 | 0.0075 | -4.7014 | 2.6076E-06 | 0.0133 |
| Superior longitudinal fasciculus | -0.0392 | 0.0077 | -5.1143 | 3.1895E-07 | 0.0028 |
| **MD** |  |  |  |  |  |
| Global MD | 0.0326 | 0.0075 | 4.3299 | 1.5015E-05 | 0.0277 |
| Acoustic radiation | 0.0339 | 0.0069 | 4.8778 | 1.0844E-06 | 0.0060 |
| Cingulate gyrus | 0.0328 | 0.0073 | 4.4648 | 8.074E-06 | 0.0248 |
| Forceps minor | 0.0348 | 0.0076 | 4.5604 | 5.1479E-06 | 0.0188 |

**Table 3.** Significant associations between SEC14L4_eQTL_cis and FA and MD-measured white matter tracts.

| **White Matter Tracts** | **Effect size** | **SD** | **t value** | **p value** | **p value, FDR corrected** |
| --- | --- | --- | --- | --- | --- |
| **FA** |  |  |  |  |  |
| Projection fibres | 0.0339 | 0.0079 | 4.3032 | 1.6943E-05 | 0.0273 |
| Forceps minor | -0.0462 | 0.0078 | -5.8981 | 3.7587E-09 | 0.0001 |
| **MD** |  |  |  |  |  |
| Forceps minor | 0.0353 | 0.0076 | 4.6349 | 3.6022E-06 | 0.0155 |

**Table 4.** Significant associations between SRA1_eQTL_cis and FA and MD-measured white matter tracts.

| **White Matter Tracts** | **Effect size** | **SD** | **t value** | **p value** | **p value, FDR corrected** |
| --- | --- | --- | --- | --- | --- |
| **FA** |  |  |  |  |  |
| Anterior thalamic radiations | 0.0324 | 0.0076 | 4.2863 | 1.8287E-05 | 0.0429 |
| Forceps minor | 0.0352 | 0.0078 | 4.4956 | 6.992E-06 | 0.0271 |
| **MD** |  |  |  |  |  |
| Global MD | -0.0328 | 0.0075 | -4.3626 | 1.2941E-05 | 0.0257 |
| Anterior thalamic radiations | -0.0339 | 0.0070 | -4.8703 | 1.1263E-06 | 0.0060 |
| Forceps minor | -0.0392 | 0.0076 | -5.1537 | 2.5879E-07 | 0.0025 |
| Inferior fronto-occipital fasciculus | -0.0335 | 0.0075 | -4.4845 | 7.3652E-06 | 0.0234 |
| Inferior longitudinal fasciculus | -0.0311 | 0.0073 | -4.2695 | 1.9718E-05 | 0.0447 |
| Superior longitudinal fasciculus | -0.0343 | 0.0076 | -4.5355 | 5.7939E-06 | 0.0204 |

**Table 5.** Significant associations between NMT1_eQTL_cis and FA and MD-measured white matter tracts.

| **White Matter Tracts** | **Effect size** | **SD** | **t value** | **p value** | **p value, FDR corrected** |
| --- | --- | --- | --- | --- | --- |
| **FA** |  |  |  |  |  |
| Forceps major | 0.0436 | 0.0081 | 5.3818 | 7.4908E-08 | 0.0009 |
| Forceps minor | 0.0338 | 0.0078 | 4.3185 | 1.5817E-05 | 0.0416 |
| **MD** |  |  |  |  |  |
| Global MD | -0.0366 | 0.0075 | -4.8650 | 1.1564E-06 | 0.0050 |
| Association fibres | -0.0368 | 0.0077 | -4.7868 | 1.7111E-06 | 0.0063 |
| Inferior longitudinal fasciculus | -0.0309 | 0.0073 | -4.2303 | 2.3485E-05 | 0.0497 |
| Superior longitudinal fasciculus | -0.0356 | 0.0076 | -4.7055 | 2.5555E-06 | 0.0116 |

**Table 6.** Significant associations between CPNE1_eQTL_cis and FA and MD-measured white matter tracts.

| **White Matter Tracts** | **Effect size** | **SD** | **t value** | **p value** | **p value, FDR corrected** |
| --- | --- | --- | --- | --- | --- |
| **FA** |  |  |  |  |  |
| Forceps minor | -0.0347 | 0.0078 | -4.4321 | 9.4015E-06 | 0.0337 |
| **MD** |  |  |  |  |  |
| Global MD | 0.0330 | 0.0075 | 4.3859 | 1.1631E-05 | 0.0250 |
| Association fibres | 0.0318 | 0.0077 | 4.1395 | 3.5002E-05 | 0.0423 |
| Thalamic radiations | 0.0296 | 0.0072 | 4.1282 | 3.6762E-05 | 0.0423 |
| Anterior thalamic radiations | 0.0356 | 0.0070 | 5.1101 | 3.2604E-07 | 0.0028 |
| Forceps minor | 0.0334 | 0.0076 | 4.3876 | 1.154E-05 | 0.0297 |
| Superior longitudinal fasciculus | 0.0342 | 0.0076 | 4.5223 | 6.1651E-06 | 0.0210 |

**Table 7.** Significant associations between PLEKHM1_eQTL_cis and FA and MD-measured white matter tracts.

| **White Matter Tracts** | **Effect size** | **SD** | **t value** | **p value** | **p value, FDR corrected** |
| --- | --- | --- | --- | --- | --- |
| **FA** |  |  |  |  |  |
| Forceps minor | -0.0382 | 0.0078 | -4.8721 | 1.1158E-06 | 0.0077 |
| **MD** |  |  |  |  |  |
| Forceps minor | 0.0331 | 0.0076 | 4.3465 | 1.3925E-05 | 0.0349 |
| Inferior fronto-occipital fasciculus | 0.0332 | 0.0075 | 4.4413 | 9.01E-06 | 0.0268 |

**Table 8.** Significant associations between UBE3C_eQTL_cis and FA and MD-measured white matter tracts.

4. **GWAS quality check and parameters**

In order to determine whether any score SNPs were previously associated with global and tract category measures of interest (i.e. tract categories & global measures found to be significantly associated with the 8 eQTL scores), we ran 8 GWAS locally (the 3 tract categories: association fibres, thalamic radiations, and projection fibres for both FA and MD, and global measures for FA and MD). We used BGENIE (1) to conduct the association analysis and excluded related participants (up to the third degree using the KING toolset (2)), as well as those who also participated in Generation Scotland and PGC MDD GWAS. Only variants with a minor allele frequency (MAF) > 0.001 (0.1%), SNP information score (quality of imputation) > 0.1, and Hardy-Weinberg equilibrium (HWE) p-value >= 1e-6 were examined. Sex, age, the first 8 principal components, genotyping array, and three head position coordinates were fitted as covariates in the analysis.

The output summary statistics files contain information with regards to the chromosome, SNP ID, p-value and effect size of association with each phenotype. We looked up the SNPs significantly associated with the tracts of interest, and noted the effect size and p-value for each.

5. **Individual white matter tracts and tract category to which they belong**

**Fractional anisotropy and mean diffusivity**

*Association fibres*

Inferior fronto-occipital fasciculus

Uncinate fasciculus

Parahippocampal cingulum

Cingulate gyrus

Superior longitudinal fasciculus

Inferior longitudinal fasciculus

*Thalamic radiations*

Anterior thalamic radiation

Posterior thalamic radiation

Superior thalamic radiation

*Projection fibres*

Acoustic radiation

Medial lemniscus

Forceps major*

Forceps minor*

Middle cerebellar peduncle*

Corticospinal tract

*Global FA & global MD*

**Table 1.** White matter tracts, global white matter and tract categories for FA and MD.

* indicates unilateral regions.

6. **P-values and effect size of each SNP for each individual white matter tract results (Elliott et al., 2018) and global and regional results (run locally); figure produced locally**

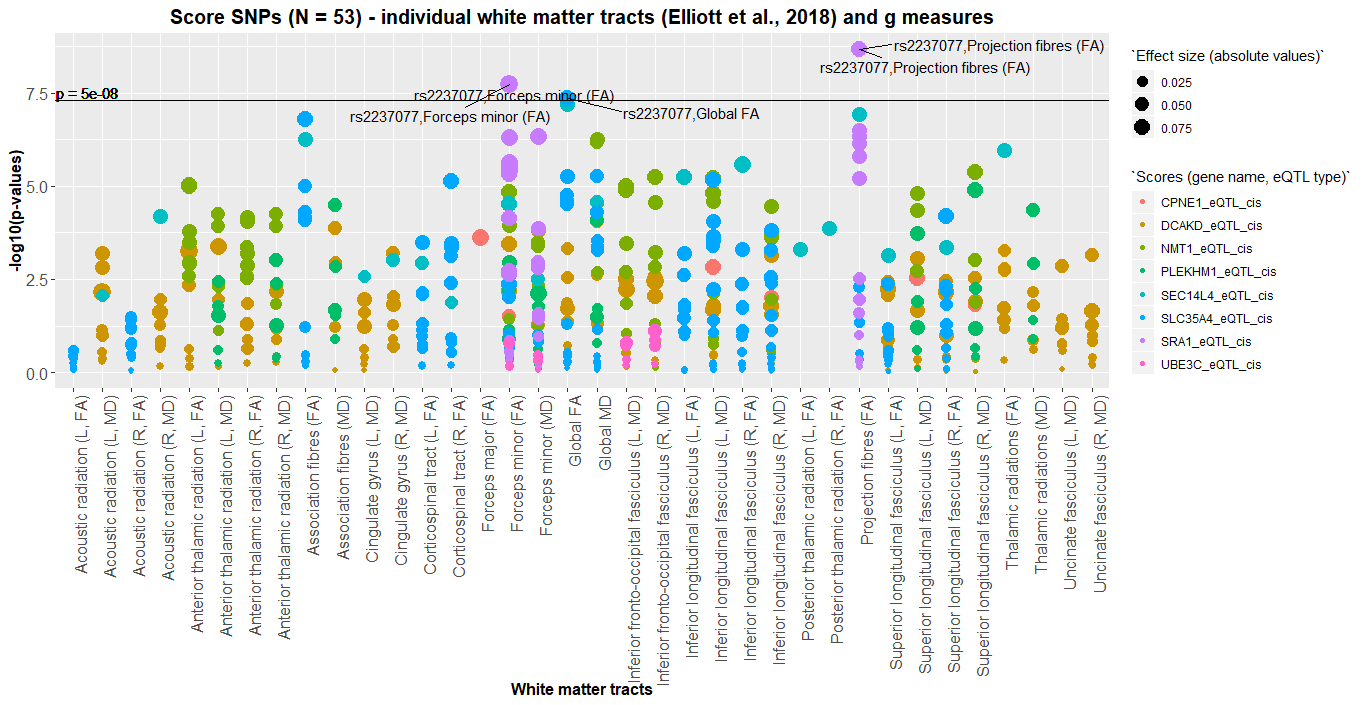

**Figure 1.** GWA between SNPs, individual white matter tracts of interest (Elliott et al., 2018) and global and tract category measures (run locally). Each point on the plot corresponds to one SNP. White matter tracts of interest are noted on the x-axis (L = left; R = right; FA = fractional anisotropy; MD = mean diffusivity). The 8 colours of the plot points indicate the score to which they belong (shown in “Scores (gene name, eQTL type)” legend). Magnitude of effect is shown in the legend entitled “Effect size (absolute values)”. The horizontal line indicates genome-wide significance (5e-8).

7. **P-values and effect size for each SNP in association with gene expression as taken from GENOSCORES; these values were obtained in the two discovery datasets used in the current study (Gusev et al., 2016; Westra et al., 2013).**

| **Chromosome** | **SNP** | **Gene** | **Effect size** | **P-value** |
| --- | --- | --- | --- | --- |
| 17 | rs4793119 | DCAKD | 0.188 | 3.98E-08 |
| 17 | rs17682536 | DCAKD | -0.114 | 1.20E-13 |
| 17 | rs962888 | DCAKD | 0.284 | 4.14E-19 |
| 17 | rs9898793 | DCAKD | 0.343 | 3.53E-240 |
| 17 | rs2040558 | DCAKD | -0.171 | 1.73E-52 |
| 17 | rs2239921 | DCAKD | 0.367 | 1.03E-28 |
| 17 | rs3744760 | DCAKD | 0.365 | 2.47E-184 |
| 17 | rs4986172 | DCAKD | 0.093 | 1.07E-24 |
| 5 | rs269783 | SLC35A4 | -0.114 | 6.55E-11 |
| 5 | rs13175916 | SLC35A4 | -0.025 | 1.33E-09 |
| 5 | rs2237077 | SLC35A4 | 0.322 | 0 |
| 5 | rs1862176 | SLC35A4 | 0.223 | 0 |
| 5 | rs6860077 | SLC35A4 | 0.210 | 0 |
| 5 | rs17286676 | SLC35A4 | -0.041 | 9.04E-97 |
| 5 | rs250430 | SLC35A4 | 0.087 | 4.09E-86 |
| 5 | rs250429 | SLC35A4 | 0.208 | 0 |
| 5 | rs12517200 | SLC35A4 | 0.061 | 7.19E-298 |
| 5 | rs1583005 | SLC35A4 | 0.138 | 5.452E-06 |
| 5 | rs2286394 | SLC35A4 | -0.055 | 4.40E-39 |
| 5 | rs3733709 | SLC35A4 | -0.110 | 5.77E-24 |
| 22 | rs2267161 | SEC14L4 | -0.093 | 6.96E-06 |
| 5 | rs2237077 | SRA1 | -0.025 | 4.55E-19 |
| 5 | rs1862176 | SRA1 | -0.040 | 1.72E-39 |
| 5 | rs6860077 | SRA1 | -0.039 | 3.91E-41 |
| 5 | rs1835959 | SRA1 | -0.087 | 1.14E-20 |
| 5 | rs250430 | SRA1 | -0.087 | 1.12E-20 |
| 5 | rs250429 | SRA1 | -0.039 | 2.05E-42 |
| 5 | rs2569163 | SRA1 | 0.048 | 3.13E-24 |
| 5 | rs778582 | SRA1 | -0.016 | 2.08E-34 |
| 5 | rs12517200 | SRA1 | -0.013 | 6.77E-34 |
| 5 | rs1583005 | SRA1 | -0.005 | 1.10E-11 |
| 5 | rs2530241 | SRA1 | 0.001 | 3.92E-14 |
| 5 | rs801186 | SRA1 | 0.010 | 3.28E-18 |
| 5 | rs801171 | SRA1 | 0.001 | 5.85E-15 |
| 5 | rs2531360 | SRA1 | 0.000 | 6.17E-14 |
| 5 | rs2240696 | SRA1 | -0.007 | 6.31E-12 |
| 17 | rs9898793 | NMT1 | 0.015 | 3.69E-08 |
| 17 | rs4793172 | NMT1 | 0.029 | 1.41E-10 |
| 17 | rs2239916 | NMT1 | 0.035 | 1.17E-14 |
| 17 | rs1053739 | NMT1 | 0.032 | 2.77E-13 |
| 17 | rs3744760 | NMT1 | 0.032 | 2.85E-10 |
| 17 | rs12946454 | NMT1 | 0.028 | 8.33E-09 |
| 17 | rs4986172 | NMT1 | 0.030 | 5.50E-09 |
| 6 | rs4324798 | CPNE1 | 0.181 | 1.79E-06 |
| 17 | rs9898793 | PLEKHM1 | 0.394 | 6.38E-78 |
| 17 | rs2239921 | PLEKHM1 | 0.304 | 6.23E-13 |
| 17 | rs3744760 | PLEKHM1 | 0.247 | 2.54E-79 |
| 17 | rs4986172 | PLEKHM1 | -0.138 | 3.73E-06 |
| 17 | rs1552458 | PLEKHM1 | 0.317 | 2.91E-47 |
| 7 | rs17646960 | UBE3C | 0.342 | 1.49E-20 |
| 7 | rs1182398 | UBE3C | -0.285 | 1.07E-33 |
| 7 | rs1182393 | UBE3C | -0.298 | 9.37E-43 |
| 7 | rs2527866 | UBE3C | -0.297 | 2.95E-06 |

**Table 1**. Associations between SNPs found in the 8 eQTL scores and gene expression (N_total_ = 53); effect size and p-values are taken from the two GWAS studies (Gusev et al., 2016; Westra et al., 2013) available in GENOSCORES.

**Supplementary material references**

1. Phillips, A. R., Suttangkakul, A., & Vierstra, R. D. (2008). The ATG12-conjugating enzyme ATG10 is essential for autophagic vesicle formation in Arabidopsis thaliana. *Genetics*, *178*(3), 1339-1353.
2. Sharma, S., Wongpalee, S. P., Vashisht, A., Wohlschlegel, J. A., & Black, D. L. (2014). Stem–loop 4 of U1 snRNA is essential for splicing and interacts with the U2 snRNP-specific SF3A1 protein during spliceosome assembly. *Genes & development*, *28*(22), 2518-2531.
3. Bansbach, C. E., Bétous, R., Lovejoy, C. A., Glick, G. G., & Cortez, D. (2009). The annealing helicase SMARCAL1 maintains genome integrity at stalled replication forks. *Genes & development*, *23*(20), 2405-2414.
4. Christopher, L., Napolioni, V., Khan, R. R., Han, S. S., Greicius, M. D., & Alzheimer's Disease Neuroimaging Initiative. (2017). A variant in PPP4R3A protects against alzheimer‐related metabolic decline. *Annals of neurology*, *82*(6), 900-911.
5. Wright, S. D., Ramos, R. A., Tobias, P. S., Ulevitch, R. J., & Mathison, J. C. (1990). CD14, a receptor for complexes of lipopolysaccharide (LPS) and LPS binding protein. *Science*, *249*(4975), 1431-1433.
6. Steet, R., & Kornfeld, S. (2006). COG-7-deficient human fibroblasts exhibit altered recycling of Golgi proteins. *Molecular biology of the cell*, *17*(5), 2312-2321.
7. Morava, E., Zeevaert, R., Korsch, E., Huijben, K., Wopereis, S., Matthijs, G., ... & Wevers, R. A. (2007). A common mutation in the COG7 gene with a consistent phenotype including microcephaly, adducted thumbs, growth retardation, VSD and episodes of hyperthermia. *European Journal of Human Genetics*, *15*(6), 638.
8. Qin, Z., Wang, Y., Tang, J., Zhang, L., Li, R., Xue, J., ... & Yang, J. (2018). High LINC01605 expression predicts poor prognosis and promotes tumor progression via up-regulation of MMP9 in bladder cancer. *Bioscience reports*, *38*(5), BSR20180562.
9. Massé, K. L., Collins, R., Bhamra, S., Seville, R. A., & Jones, E. (2007). Anxa4 genes are expressed in distinct organ systems in xenopus laevis and tropicalis but are functionally conserved. *Organogenesis*, *3*(2), 83-92.
10. Ergin, V., Erdogan, M., & Menevse, A. (2015). Regulation of shootin1 gene expression involves ngf-induced alternative splicing during neuronal differentiation of PC12 cells. *Scientific reports*, *5*, 17931.
11. Yeo, S. Y., Ha, S. Y., Yu, E. J., Lee, K. W., Kim, J. H., & Kim, S. H. (2014). ZNF282 (Zinc finger protein 282), a novel E2F1 co-activator, promotes esophageal squamous cell carcinoma. *Oncotarget*, *5*(23), 12260.
12. Edqvist, P. H. D., Huvila, J., Forsström, B., Talve, L., Carpén, O., Salvesen, H. B., ... & Uhlén, M. (2015). Loss of ASRGL1 expression is an independent biomarker for disease-specific survival in endometrioid endometrial carcinoma. *Gynecologic oncology*, *137*(3), 529-537.
13. Bhattacharya, M. R., Geisler, S., Pittman, S. K., Doan, R. A., Weihl, C. C., Milbrandt, J., & DiAntonio, A. (2016). TMEM184b promotes axon degeneration and neuromuscular junction maintenance. *Journal of Neuroscience*, *36*(17), 4681-4689.
14. Wang, L., Wildt, K. F., Castro, E., Xiong, Y., Feigenbaum, L., Tessarollo, L., & Bosselut, R. (2008). The zinc finger transcription factor Zbtb7b represses CD8-lineage gene expression in peripheral CD4+ T cells. *Immunity*, *29*(6), 876-887.
15. Nesbit, M. A., Hannan, F. M., Howles, S. A., Reed, A. A., Cranston, T., Thakker, C. E., ... & Morrison, P. J. (2013). Mutations in AP2S1 cause familial hypocalciuric hypercalcemia type 3. *Nature genetics*, *45*(1), 93.
16. Spiegel, R., Shaag, A., Shalev, S., & Elpeleg, O. (2016). Homozygous mutation in the APOA1BP is associated with a lethal infantile leukoencephalopathy. *Neurogenetics*, *17*(3), 187-190.
17. Zhu, M., Yan, C., Ren, C., Huang, X., Zhu, X., Gu, H., ... & Miao, X. (2017). Exome array analysis identifies variants in SPOCD1 and BTN3A2 that affect risk for gastric cancer. *Gastroenterology*, *152*(8), 2011-2021.
18. Evans, D. R., & Guy, H. I. (2004). Mammalian pyrimidine biosynthesis: fresh insights into an ancient pathway. *Journal of Biological Chemistry*, *279*(32), 33035-33038.
19. Maxson, J. E., Gotlib, J., Pollyea, D. A., Fleischman, A. G., Agarwal, A., Eide, C. A., ... & Pond, J. B. (2013). Oncogenic CSF3R mutations in chronic neutrophilic leukemia and atypical CML. *New England Journal of Medicine*, *368*(19), 1781-1790.
20. Rhee, H. W., Zou, P., Udeshi, N. D., Martell, J. D., Mootha, V. K., Carr, S. A., & Ting, A. Y. (2013). Proteomic mapping of mitochondria in living cells via spatially restricted enzymatic tagging. *Science*, *339*(6125), 1328-1331.
21. Zollino, M., Orteschi, D., Murdolo, M., Lattante, S., Battaglia, D., Stefanini, C., ... & Marangi, G. (2012). Mutations in KANSL1 cause the 17q21. 31 microdeletion syndrome phenotype. *Nature genetics*, *44*(6), 636.
22. Zhernakova, A., Festen, E. M., Franke, L., Trynka, G., van Diemen, C. C., Monsuur, A. J., ... & Boezen, H. M. (2008). Genetic analysis of innate immunity in Crohn's disease and ulcerative colitis identifies two susceptibility loci harboring CARD9 and IL18RAP. *The American Journal of Human Genetics*, *82*(5), 1202-1210.
23. Gertow, K., Sennblad, B., Strawbridge, R. J., Öhrvik, J., Zabaneh, D., Shah, S., ... & Kivimäki, M. (2012). Identification of the BCAR1-CFDP1-TMEM170A locus as a determinant of carotid intima-media thickness and coronary artery disease risk. *Circulation: Cardiovascular Genetics*, *5*(6), 656-665.
24. Zhu, J. H., Chen, R., Yi, W., Cantin, G. T., Fearns, C., Yang, Y., ... & Lee, J. D. (2008). Protein tyrosine phosphatase PTPN13 negatively regulates Her2/ErbB2 malignant signaling. *Oncogene*, *27*(18), 2525.
25. Wills, Z., Bateman, J., Korey, C. A., Comer, A., & Van Vactor, D. (1999). The tyrosine kinase Abl and its substrate enabled collaborate with the receptor phosphatase Dlar to control motor axon guidance. *Neuron*, *22*(2), 301-312.
26. Niwa, T., Saito, H., Imajoh‐ohmi, S., Kaminishi, M., Seto, Y., Miki, Y., & Nakanishi, A. (2009). BRCA2 interacts with the cytoskeletal linker protein plectin to form a complex controlling centrosome localization. *Cancer science*, *100*(11), 2115-2125.
27. Bycroft, C., Freeman, C., Petkova, D., Band, G., Elliott, L. T., Sharp, K., ... & Cortes, A. (2018). The UK Biobank resource with deep phenotyping and genomic data. Nature, 562(7726), 203.
28. Manichaikul, A., Mychaleckyj, J. C., Rich, S. S., Daly, K., Sale, M., & Chen, W. M. (2010). Robust relationship inference in genome-wide association studies. Bioinformatics, 26(22), 2867-2873.
